# Structural and functional characterization of DdrC, a novel DNA damage-induced nucleoid associated protein involved in DNA compaction

**DOI:** 10.1101/2021.10.27.466113

**Authors:** Anne-Sophie Banneville, Claire Bouthier de la Tour, Cécilia Hognon, Jacques-Philippe Colletier, Jean-Marie Teulon, Aline Le Roy, Jean-Luc Pellequer, Antonio Monari, François Dehez, Fabrice Confalonieri, Pascale Servant, Joanna Timmins

## Abstract

*Deinococcus radiodurans* is a spherical bacterium well-known for its outstanding resistance to DNA-damaging agents. Exposure to such agents leads to drastic changes in the transcriptome of *D. radiodurans*. In particular, four *Deinococcus*-specific genes, known as DNA Damage Response genes, are strongly up-regulated and have been shown to contribute to the resistance phenotype of *D. radiodurans*. One of these, DdrC, is expressed shortly after exposure to γ-radiation and is rapidly recruited to the nucleoid. *In vitro*, DdrC has been shown to compact circular DNA, circularize linear DNA, anneal complementary DNA strands and protect DNA from nucleases. To shed light on the possible functions of DdrC in *D. radiodurans*, we determined the crystal structure of the domain-swapped DdrC dimer at a resolution of 2.2 Å and further characterized its DNA binding and compaction properties. Notably, we show that DdrC bears two asymmetric DNA binding sites located on either side of the dimer and can modulate the topology and level of compaction of circular DNA. These findings suggest that DdrC may be a DNA damage-induced nucleoid-associated protein that enhances nucleoid compaction to limit the dispersion of the fragmented genome and facilitate DNA repair after exposure to severe DNA damaging conditions.

## INTRODUCTION

*Deinococcus radiodurans* is a gram-positive spherical bacterium, highly resistant to DNA-damaging agents including ionizing radiation, UV-light, desiccation and reactive oxygen species (1). Several mechanisms have been proposed to contribute to the maintenance of proteome and genome integrity in this bacterium: (i) multiple anti-oxidant strategies including a high intracellular Mn/Fe ratio (2), (ii) efficient DNA repair systems (3, 4) and (iii) a highly compact nucleoid (5–7), which may limit dispersion of DNA fragments, thus easing DNA repair processes. Following exposure to ionizing radiation or desiccation, the transcriptome of *D. radiodurans* undergoes a drastic change with the up-regulation of many genes, several of which are involved in DNA repair (8). Interestingly, in *D. radiodurans*, this up-regulation has been shown to involve an SOS-independent response system, the radiation-desiccation response (RDR) regulon that is negatively regulated at the transcriptional level under normal growth conditions by the transcriptional repressor, DdrO (9–13). DdrO binds to a conserved 17 bp palindromic sequence named the radiation-desiccation response motif (RDRM), located in the promoter regions of regulated genes (10, 14–16). Exposure of cells to radiation or desiccation leads to the activation of IrrE, a constitutively expressed metalloprotease, that cleaves DdrO leading to its inactivation, thereby relieving its negative control over gene expression (10, 16–20).

After exposure to radiation or desiccation, four of the most up-regulated genes in the RDR regulon are *ddrA, ddrB, ddrC* and *ddrD* (21). These DNA damage response proteins (Ddr) are DNA-binding proteins specific to *Deinococcus* species. After exposure to ionizing radiation, these proteins are rapidly recruited to the nucleoid where they perform distinct functions (8, 22). DdrA preferentially binds *in vitro* to 3’ single-stranded DNA (ssDNA) ends, protecting them from degradation by exonucleases and has thus been proposed to be part of an end-protection system (23, 24). DdrB is an ssDNA-binding (SSB)-like protein that promotes single-strand annealing (SSA), thereby playing an important role in the assembly of small chromosomal fragments produced by exposure to high doses of γ-radiation (25, 26). DdrB is also involved in plasmid transformation, through its SSA activity that enables the reconstitution of double-stranded DNA (dsDNA) plasmid after its internalization (26, 27). Recent studies have showed that DdrD is a ssDNA binding protein that likely contributes to genome reconstitution following exposure to irradiation (28).

DdrC is a 25 kDa DNA-binding protein that is highly overexpressed shortly after irradiation and is rapidly recruited to the nucleoid, where it has been proposed to interact with damaged DNA (8, 29). Interestingly, DdrC is distributed all over the nucleoid shortly after irradiation, but after 2 to 3 hours, it forms discrete foci located at the sites of septal closure in between the newly segregated chromosomes of *D. radiodurans* (29). *In vitro* assays have shown that DdrC binds both ss- and dsDNA, with a preference for ssDNA. This protein exerts many activities upon DNA-binding, such as compaction of circular DNA, circularization of linear DNA, annealing of complementary DNA strands and protection of DNA from nucleases. These pleiotropic activities suggest that DdrC may play a role in the repair of radiation-induced DNA damage by preventing the dispersion of DNA fragments and participating in single-strand annealing.

To shed further light on the possible functions of *D. radiodurans* DdrC in the response to DNA damage, we here focused on elucidating its three-dimensional crystal structure and on characterizing its DNA binding properties using a combination of biochemical, biophysical and computational approaches. In the absence of any structures of DdrC homologues, we solved its structure *de novo* by use of the single-wavelength anomalous dispersion method (SAD). The structure reveals that DdrC is a largely α-helical protein, composed of an N-terminal winged helix-turn-helix (wHTH) motif and a C-terminal four-helix bundle, that folds as a domain-swapped dimer. We reveal that DdrC possesses two asymmetric DNA binding sites, one on either side of the dimer formed by motifs from both its N- and C-terminal domains. We also demonstrate that DdrC can modify the topology and induce a strong compaction of circular plasmid DNA in a concentration-dependent manner. Together these findings indicate that DdrC may be a novel DNA damage-induced nucleoid-associated protein (NAP) that is recruited to the nucleoid in response to irradiation to modulate the extent of compaction of the genome and facilitate DNA repair processes.

## MATERIAL AND METHODS

### Expression and purification of DdrC and DdrC-SeMet

The *ddrC* gene (A2G07_003810) was amplified from *D. radiodurans* genomic DNA by PCR and cloned into pProExHtB (EMBL) expression vector for expression with a cleavable N-terminal His-tag. DdrC was expressed in *E. coli* BL21(DE3) cells grown in LB supplemented with 100 μg ml^−1^ ampicillin. Expression was induced with 1 mM IPTG at 28°C for 4 hours. Cells were pelleted by centrifugation and resuspended in 40 ml lysis buffer composed of 50 mM Tris-HCl pH 7.5, 0.8 M NaCl, 5 mM MgCl_2_, 10% (w/v) sucrose, 0.01% (v/v) triton X-100, 1 μg ml^−1^ DNaseI, 1 μg ml^−1^ lysozyme and a tablet of complete EDTA-free Protease Inhibitor Cocktail (Roche). Cells were lysed by sonication on ice for 3 min and centrifuged at 48,300 × g for 30 min. The cleared supernatant was loaded on a 5 ml HisTrap FF nickel affinity column (GE Healthcare), pre-equilibrated with buffer A (50 mM Tris-HCl pH 7.5, 0.8 M NaCl, 1 mM MgCl_2_). After washing the column with buffer A, buffer A supplemented with 25 mM imidazole and buffer A supplemented with 50 mM imidazole, DdrC was eluted with a linear gradient of imidazole from 50 to 500 mM imidazole in buffer A. The fractions containing DdrC were pooled and dialyzed overnight at 4°C against buffer A supplemented with 5% (v/v) glycerol in the presence of 1:20 (w/w) TEV protease to cleave the His-tag. The His-tag itself and traces of uncleaved protein were subsequently removed by nickel affinity chromatography on 1 ml Ni-Sepharose 6 FF resin (GE Healthcare) pre-equilibrated in buffer A. The cleaved DdrC was recovered in the flow-through and in a 25 mM imidazole wash, and was concentrated prior to size exclusion chromatography on a Superdex 75 10/300 GL column (GE Healthcare) pre-equilibrated with buffer B (20 mM Tris-HCl pH 7.5, 200 mM NaCl, 5% (v/v) glycerol). Finally, DdrC was concentrated to a final concentration of 24 mg ml^−1^ and stored at −80°C. For the AFM experiments, a batch of DdrC was produced following the protocol described previously, but without glycerol in the purification buffers. This ‘glycerol-free’ batch was stored at a concentration of 16 mg ml^−1^ at −80° C. The selenomethionine substituted DdrC (SeMet-DdrC) was produced in *E. coli* BL21(DE3) cells grown at 37°C in minimal M9 medium supplemented with 100 μg ml^−1^ ampicillin using a modified version of the metabolic inhibition protocol described previously (30, 31). Expression was induced overnight with 1 mM IPTG at 28°C. The SeMet-DdrC protein was then purified as described for native DdrC and was stored at 20 mg ml^−1^ in buffer B at −80°C.

### Crystallization of DdrC and SeMet-DdrC

Initial crystallization hits were obtained by robotic screening at the HTX lab (EMBL) using nanoliter sitting drops at 20°C (32). Crystals grew after 2 to 3 months in conditions containing 1.6 M ammonium sulfate and 0.1 M Tris pH 8.0 or Bicine pH 9.0. Manual crystallization screens were then performed using the hanging-drop vapor-diffusion method in 24-well plates at 20°C. Briefly, 1 μl protein solution (at 24 mg ml^−1^ for native DdrC or 20 mg ml^−1^ for SeMet-DdrC) was mixed with 1 μl mother liquor solution and equilibrated against 500 μl mother liquor solution. Crystallization conditions were refined using 0.1 M Tris pH 8.0 to pH 8.5 or Bicine pH 9.0 to pH 9.5 and 1.0 M to 2.1 M ammonium sulfate. Hexagonal bipyramidal or triangular prism-shaped crystals of DdrC and SeMet-DdrC appeared after 3-4 weeks in all conditions with ammonium sulfate below 1.9 M. Crystals were transferred to mother liquor containing 20% (v/v) glycerol as a cryoprotectant and flash-cooled in liquid nitrogen before data collection. The best diffracting crystals were obtained in 0.1 M Tris pH 8.0, 1.2 M ammonium sulfate for native DdrC and 0.1 M Tris pH 8.5, 1.9 M ammonium sulfate for SeMet-DdrC.

### Data collection and structure determination

A selenium single-wavelength anomalous diffraction (Se-SAD) dataset was collected on a SeMet-DdrC crystal at 100 K on beamline ID23-1 at the European Synchrotron Radiation Facility (Grenoble, France), on a Pilatus 6M detector (Dectris). A total of 500 images were collected at a wavelength of 0.978 Å with 100 ms exposure and an oscillation angle of 0.15° per frame. For the native DdrC, data collection was performed at 100 K on beamline Proxima-2A at the SOLEIL (Paris, France), on a Eiger 9 M detector (Dectris). A total of 3600 images were collected at a wavelength of 0.980 Å with 25 ms exposure and an oscillation angle of 0.1° per frame. Data were integrated, indexed and scaled with XDS (33). The space group was P3_2_21 for both the SeMet variant and the native DdrC, with unit cell parameters of a = 111.36 Å, b = 111.36 Å, c = 104.88 Å and a = 111.28 Å, b = 111.28 Å, c = 104.71 Å respectively (Table 1). The Se-SAD dataset was processed with the CRANK2 suite (34). Briefly, SHELXC (35) was used to calculate structure factor estimates from merged intensities, after what heavy-atom search was performed using SHELXD (35) and a resolution cutoff of 3.69 Å. Two selenium sites were found. The substructure refinement and phasing were performed with REFMAC5 (36), then experimental phases were improved using density modification with PARROT (37), which enabled automatic determination of the correct hand. Automatic model building and structure refinement were performed with Buccaneer (38, 39) and REFMAC5. Refinement converged at an R_work_ and R_free_ of 34.3 % and 40.0 %, respectively. The native DdrC structure was solved by molecular replacement with Phaser MR (40) using the SeMet-DdrC structure as a search model to a resolution of 2.2 Å (Table 1). The DdrC structure was then refined using iterative cycles of manual building in Coot (41) and refinement in PHENIX (42). The final R_work_ and R_free_ are 26.6 % and 30.3 % respectively. The structure of DdrC was validated in MolProbity (43) with only 3 residues in the outliers region of the Ramachandran plot (Table 1) and deposited in the Protein Data Bank with accession number ####. Analysis of dimerization interface and crystal contacts was carried out using PISA (44). Electrostatic surfaces (at pH 7.5 and 200 mM NaCl) were produced by APBS (45) from structures protonated by PDB2PQR (46) after structure-based titration of protonatable residues using PROPKA (47).

**Table 1.**
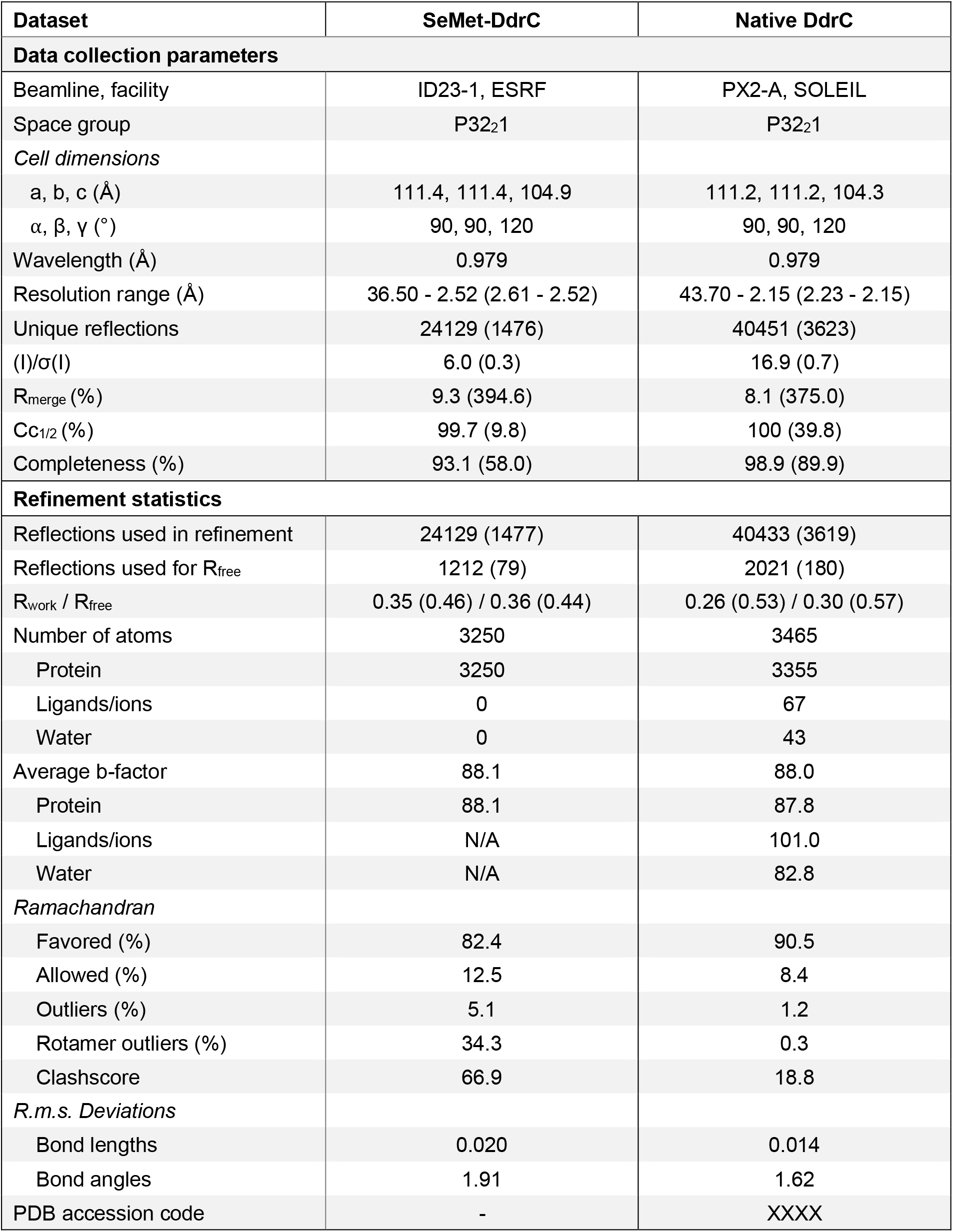
Crystallographic data collection and refinement statistics. Values in parentheses are for the highest resolution shell.

### DdrC structure prediction

The sequence of DdrC was submitted to AlphaFold2 (48) via the Colaboratory service from Google Research (https://colab.research.google.com/github/sokrypton/ColabFold/blob/main/beta/AlphaFold2_advanced.ipynb) and to RosettaFold (49) (https://robetta.bakerlab.org). Of note, the mmseq2 method (50, 51) was employed for the multiple-sequence alignment. The first five models predicted by each program were overlaid using the *align* tool in PyMOL (52) with overall root mean square deviation (rmsd) values of 0.312 to 0.498 Å for RosettaFold and of 0.266 to 0.735 Å for AlphaFold2. For both programs, the C-terminal domain was predicted with high accuracy, but only AlphaFold2 offered accurate prediction of the N-terminal domain (Supp. Table S1 and Fig. S1). Furthermore, AlphaFold2 was able to predict the overall fold of DdrC monomer A correctly, due to correct structure prediction for the linker segment, although the orientation of the NTD relative to the CTD was a little off (Supp. Fig. S1). Nonetheless, attempts to phase the native crystallographic data by molecular replacement with Phaser (40) using the best-ranked model from AlphaFold2 in its entirety as a search model, and looking for either one or two copies of the protein, failed. Similar results were obtained when using isolated N- and C-terminal domains as search models, asking for placement of either one or two copies of each, or just one or two copies of the more correctly predicted C-terminal domain. Regardless of the strategy, no solution was obtained that yielded R_free_/R_work_ values indicative of success after reciprocal space refinement using Refmac5 (10 cycles) (53). We furthermore subjected the top three molecular replacement solutions in each case to the buccaneer pipeline in CCP4 (38) with the aim to verify if automatic model-building and refinement would have been possible based on these, but once again this was unsuccessful. Thus, neither the prediction from AlphaFold2 nor that from RosettaFold would have allowed successful phasing by molecular replacement. Finally, we attempted to predict the dimer structure of DdrC using AlphaFold2, yet none of the predicted dimer models came close to the experimentally determined structure of DdrC, likely due to the bias induced by prediction of the same monomeric structure for the two monomers in the dimer.

### Preparation of supercoiled plasmid DNA

Plasmid pUC19 DNA was amplified in DH5α *E. coli* cells grown in LB with 100 μg ml^−1^ ampicillin. The supercoiled pUC19 (pUC19sc) was extracted from 100 ml overnight cultures using the NucleoBond Xtra Midi kit (Macherey-Nagel) following manufacturer’s instructions. The final DNA resuspension was performed in 50 μl milli-Q water, yielding pUC19sc at a concentration of 1.5 μg μl^−1^ (equivalent to 900 nM). The stock solution was stored at −20°C.

### Atomic Force Microscopy

pUC19sc was diluted in milli-Q water to a final concentration of 0.5 nM for all samples. ‘Glycerol-free’ DdrC was diluted in milli-Q water to a final concentration of either 2 nM, 5 nM, 10 nM or 20 nM. For the protein-DNA samples, pUC19sc was incubated with DdrC for 30 min at 30°C before sample deposition on the mica sheet. Topographic data were acquired by a multimode 8 microscope equipped with a Nanoscope V controller (Bruker, Santa Barbara, USA). Before use, a freshly cleaved V-1 grade muscovite mica (Nanoandmore, Wetzlar, Germany) sheet was pre-treated with 10 μl 5 mM NiCl_2_ and dried under nitrogen gas. 5 μl of each sample solution was deposited on the mica, after which the mica was incubated for 2 min, then dried under a gentle stream of nitrogen gas. All imaging was conducted with the PeakForce Tapping mode and ScanAsyst mode at a rate of ~1.0 Hz; the resolution was set to either 512 or 1024 pixels per scan line. The SCANASYST-AIRHR cantilever was employed with nominal values of k = 0.4 N m^−1^, Fq = 130 kHz and tip radius = 2 nm (Bruker probes, Camarillo, CA, USA). Whenever the ScanAsyst mode was applied, a semi-manual control was on during the imaging procedure to manually adjust the set point and gain in order to reduce the tip-sample interactions to the minimum. The ramp size was kept constant at 150 nm. Processing of raw AFM images was systematically performed using the Gwyddion software (54). First, raw AFM images were flattened using a plane fit to the first order, then the flattening effect was further enhanced by applying the “line flattening” tool of Gwyddion with a polynomial of order 3, followed by exclusion of all imaged objects whose height values exceeded the given threshold (usually 0.1 – 0.5 nm). When necessary, stripe noises were reduced using the ‘Remove Scars’ function in Gwyddion. Measurements of the surface areas of individual assemblies were performed on these processed AFM images corresponding to 2 μm^2^ or 1 μm^2^ areas. A classical height threshold was applied on the image to select as many individual assemblies as possible. Assemblies that either touched the border of the image or were not clearly identifiable due to unresolved overlapping (three or more plasmids in a single selection) were excluded from the statistical analysis. The surface areas of the selected assemblies were extracted using the grain distribution function in Gwyddion. To discriminate between the condensed or more opened assemblies, a surface area threshold of 6000 nm to 9500 nm2 was applied to each image depending on their respective height threshold used for selection. Histograms and scatter-plots representing the fraction of condensed assemblies as a function of DdrC concentration were then plotted using the GraphPad Prism 8 software.

### Fluorescence polarization

Equilibrium fluorescence polarization DNA binding assays were performed on a Clariostar (BMG Labtech) microplate reader, fitted with polarization filters. Reactions were performed in black 386-well medium-binding plates (Greiner). 0 to 400 μM DdrC (dimer) were titrated into 10 nM 5’-FAM labelled dsDNA 20 mer or 50 mer substrates (Table S1) in binding buffer composed of 20 mM Tris pH 8.0, 100 mM NaCl, 1 mM MgCl_2_ and 0.2 mg ml^−1^ BSA. Reactions were performed in a final volume of 40 μl at room temperature. After subtracting the polarization values obtained for DNA alone, the mean data from at least three independent measurements were fitted to one of the following equations using GraphPad Prism 8: (a) a simple one-site specific binding model (Y=(B_max_*X)/(K_D_+X)), (b) a one-site specific binding model with Hill coefficient (Y=(B_max_*X^h^)/(K_D_^h^+X^h^)), or (c) a two-site specific binding model (Y=[(B_max_(Hi)*X)/(K_D_(Hi)+X)]+[(B_max_(Lo)*X)/(K_D_(Lo)+X)]), where Y is the difference between the anisotropy of completely bound and completely free oligo, B_max_ is the maximal polarization signal, X is the DdrC concentration, K_D_ is the equilibrium dissociation constant and h is the Hill slope (Table S2).

### Analysis of plasmid topology by 1D and 2D gel electrophoresis

200 ng of relaxed DNA (relaxed pHOT-1 DNA, 2.4 kb) (Topogen) was incubated for 15 min at 4°C in the absence or presence of increasing concentrations of DdrC in 25 μl of buffer composed of 40 mM Tris-HCl pH7.8, 5 mM MgCl_2_, 1.5 mM DTT, 50 mM NaCl, 12% (v/v) glycerol. 10 U of topoisomerase I from wheat germ (Sigma) was then added and the samples were incubated 30 min at 30°C. Reactions were stopped by addition of a mix of 1 mg ml^−1^ Proteinase K and 0.5% (w/v) SDS followed by an incubation at 37°C for 10 min. 7 μl 6X DNA Loading Dye were then added to the reactions and 10 μl of the reaction mixtures were separated by gel electrophoresis at 4°C on 1.2% agarose gels in TEP buffer (36 mM Tris-HCl pH 7.8, 30 mM NaH_2_PO4, 1 mM EDTA) at 4.3V/cm for 4h. DNA topoisomers were revealed after ethidium bromide staining. For 2D gel electrophoresis, 20 μl of the remaining reaction mixtures were loaded on a 1.2% agarose gel. The first dimension was performed as described above. The second dimension was run in a perpendicular direction at 1V/cm for 16h at room temperature in TEP buffer containing 3 μg ml^−1^ chloroquine, a DNA intercalator that unwinds the double helix of a closed circular DNA, resulting in a loss of negative supercoils and formation of positive supercoils. The chloroquine was then eliminated from the gel by incubation in H_2_O for 3h and the distribution of topoisomers was visualized after ethidium bromide staining.

### Size-exclusion chromatography coupled to multi-angle laser light scattering

Size exclusion chromatography (SEC) combined with multi-angle laser light scattering (MALLS), dynamic light scattering (DLS) and refractometry (RI) experiments were performed with a Superdex 200 10/300 GL size exclusion column equilibrated with Buffer C (20 mM Tris pH 7.5 and 200 mM NaCl) at room temperature. 20 μl DdrC at 16 mg ml^−1^ was injected onto the column at 0.5 ml min^−1^. On-line MALLS detection was performed with a miniDAWN-TREOS detector (Wyatt), DLS was recorded with a DynaPro Nanostar and RI measurements were performed with an Optilab eEX system (Wyatt).

### Analytical ultracentrifugation

Sedimentation velocity experiments were performed at 42,000 rpm and 4°C, on a Beckman XLI analytical ultracentrifuge using a AN-60 Ti rotor (Beckman Coulter, Brea, USA) and double-sector cells with optical path lengths of 12 and 1.5 mm equipped with sapphire windows (Nanolytics, Potsdam, DE). Buffer C was used as a reference. Measurements were made on 1 mg ml^−1^, 4 mg ml^−1^ and 8 mg ml^−1^ DdrC using absorbance at 280 nm and interference optics. Data were processed with the REDATE software (https://www.utsouthwestern.edu/labs/mbr/software/) and the parameters were determined with SEDNTERP (55) and SEDFIT (56). Analysis of sedimentation coefficients and molecular weights were performed using SEDFIT and GUSSI (57).

### Molecular dynamics simulations

The domain-swapped DdrC dimer from the asymmetric unit was used as a starting model for all-atoms MD simulations after building the missing loops between helices α7 and α8 in chains A and B using the loop modelling tool in Modeller (58). Two MD simulations were performed on the apo-DdrC structure to verify the stability of the dimer and evaluate the overall dynamics of DdrC. For protein-DNA assemblies, two 25 bp DNA duplexes of random sequence were manually positioned on either side of the DdrC dimer following the positively charged grooves so as to minimize steric clashes between the DNA and protein side chains. DdrC-bound to two DNA duplexes were then used for five independent MD simulations to enhance the statistical sampling. All the macromolecular systems were explicitly hydrated in boxes of 42,000 water molecules containing 22 sodium ions to ensure the overall electrical neutrality of the unit cells. Water was represented by means of the TIP3P water model (59), whereas protein, DNA and ions were described using the amberf99 force field (60) including the bsc1 corrections for DNA (61). All setups were generated using the tleap facility of Amber Tools (62, 63). Molecular rendering and analyses were done using VMD (64). MD simulations were performed using the massively parallel code NAMD (65). All trajectories were generated in the isobaric-isothermal ensemble, at 300 K under 1 atm using Langevin dynamics (66) (damping coefficient 1 ps^−1^) and the Langevin piston method (67), respectively. Long-range electrostatic interactions were accounted for by means of the Particle Mesh Ewald (PME) algorithm (68). The rattle algorithm was used to constrain lengths of covalent bonds involving hydrogen atoms to their equilibrium value (69). The classical equations of motion were integrated through a time step of 4 fs using the hydrogen mass repartition strategy (70). Each molecular assay was thermalized during 15 ns, followed by 500 ns of production.

## RESULTS

### *De novo* phasing of the structure of DdrC - an unusual asymmetric domain-swapped dimer

DdrC is a protein for which no known structural homologues have been identified. We therefore solved the structure of *D. radiodurans* DdrC *de novo* by use of the single-wavelength anomalous dispersion method (SAD). We determined the structure of a selenomethionine variant of DdrC (SeMet-DdrC) by SAD to a resolution of 2.5 Å, and then solved the structure of native DdrC by molecular replacement using the SeMet-DdrC as a search model and refined it to 2.2 Å resolution (Table 1). The asymmetric unit contains a DdrC dimer composed of chains A and B. Almost all residues are visible in the electron density, notwithstanding 3 terminal residues missing at both the N- (residues 1 to 3) and C-termini (residues 229 to 231) and a highly disordered loop between helices α7 and α8 (Fig. 1A and B), corresponding to residues 167-173 and 161-169 in chains A and B, respectively.

**Figure 1.**
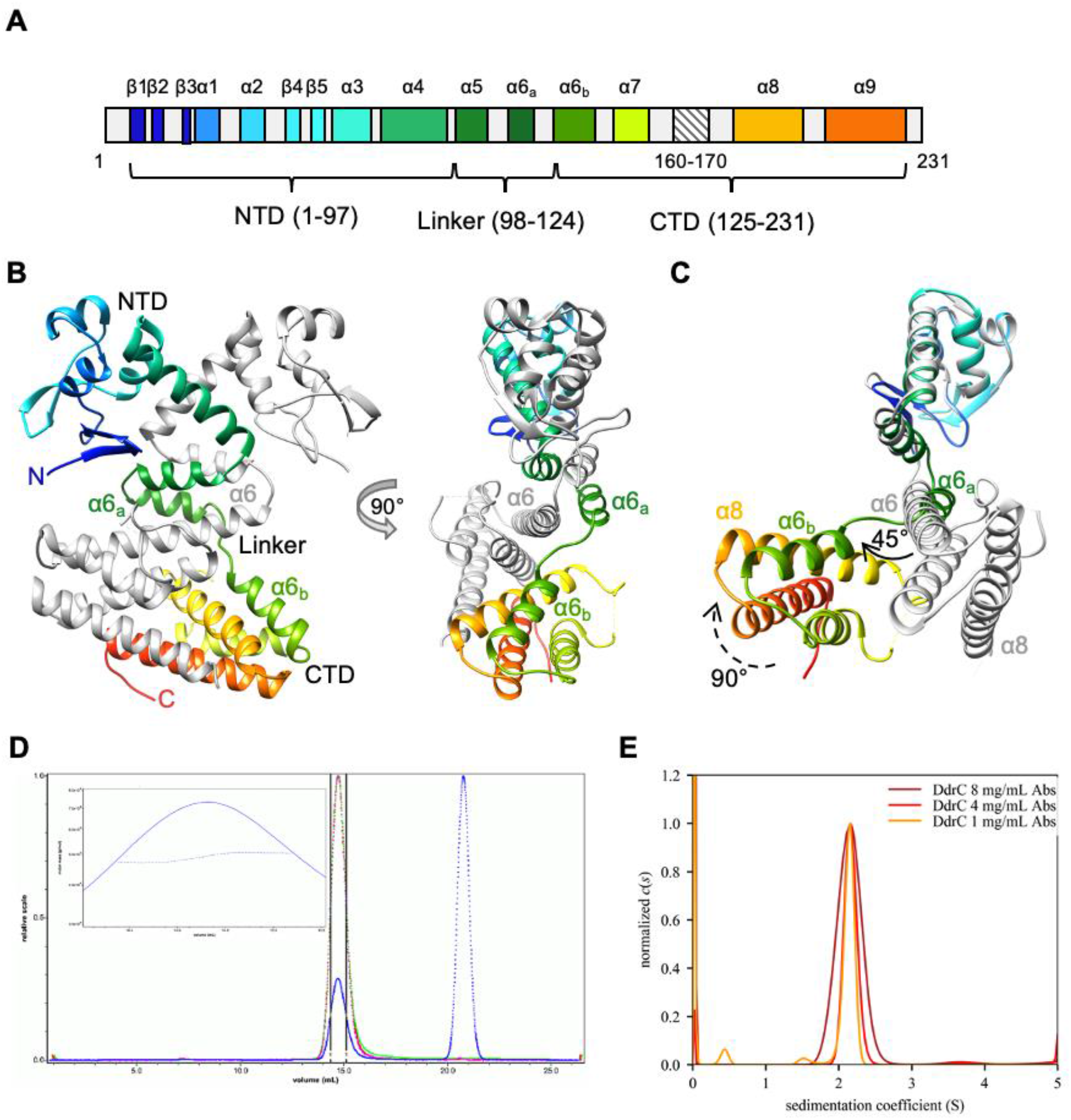
DdrC is an unusual domain-swapped dimer composed of two domains. (A) Secondary structure organization of DdrC (chain B), colored from blue (N-terminus) to orange (C-terminus). Residues 160 to 170 were not visible in the structure and are thus expected to form a disordered loop. NTD: N-terminal domain, CTD: C-terminal domain. (B) Front and side views of the DdrC dimer, with monomer A colored in gray and monomer B colored in rainbow colors from blue (N-terminus) to red (C-terminus). The side view of DdrC highlights the asymmetry between the two faces of the dimer. (C) Side view of the overlay of the two DdrC chains using the NTD as a reference. Chains A and B are colored as in (B). The α6_a_ and α6_b_ helices in chain B correspond to a distorted conformation of the long α6 helix of chain A, probably to accommodate the domain swapping of the two monomers. (D) Size-exclusion chromatogram obtained from SEC-MALLS analysis of DdrC. The inset represents a close-up view of the DdrC peak (defined by black lines), illustrating molar mass points obtained along the peak. The mean mass of DdrC derived from this data was 49.1 kDa, corresponding to a dimer. (E) Distribution of sedimentation coefficients obtained by analytical ultracentrifugation analysis of DdrC at three concentrations: 1 mg ml^−1^ (orange), 4 mg ml^−1^ (red) and 8 mg ml^−1^ (dark red). A majority of the sample (94 ± 4%) was found in a peak at 2.15S corresponding to a mean mass of 43.5 ± 3.5 kDa, corresponding here again to a dimer.

Recently, it was shown that use of artificial intelligence in programs such as AlphaFold2 (48) or RosettaFold (49) could enable prediction of protein structures to an accuracy high enough to allow phasing of crystallographic data by molecular replacement. To determine whether this would have been possible in the case of DdrC, we submitted the sequence of DdrC to the two programs and then attempted phasing of the native crystallographic data by molecular replacement using the best-ranked model from AlphaFold2 in its entirety or as isolated N- and C-terminal domains as a search model (see Materials and Methods for details; Supp. Table S1 and Fig. S1). Regardless of the strategy, no solution was obtained that yielded R_free_/R_work_ values indicative of success and when these putative molecular replacement solutions were submitted to automatic model-building and refinement programs, there again, they failed to produce a reliable solution. In the case of DdrC at least, experimental phasing thus turned out to be the only route towards structure elucidation.

The asymmetric unit contains a domain-swapped homo-dimer of DdrC, in which each monomer buries on average 3214 ± 24 Å^2^, corresponding to 36% of its surface area, within the dimer interface that is stabilized by 16 H-bonds and 16 salt bridges (44). Each monomer of DdrC is composed of two domains connected by a linker region (Fig. 1A and B). The N-terminal domain (NTD; residues 1-97) comprises five β-strands and four α-helices adopting a classic winged-helix-turn-helix (wHTH) motif (β3 to β5 and α1 to α3) preceded by a β-hairpin (β1 and β2) and followed by an α-helix (α4) that provides the first contacts for dimerization. The C-terminal domain (CTD; residues 125-231), which is domain-swapped between the two monomers, is composed of four α-helices organized in a four-helix bundle motif (α6_b_ to α9). The two domains are connected by a linker region comprising residues 98-124 that encompass helix α5 and the N-terminal region of α6 (α6_a_).

Although DdrC is homo-dimeric, there is a remarkable asymmetry between the two chains, which is rarely observed in domain-swapped dimers (Fig. 1C and Supp. Table S1). The two monomers of DdrC display distinct conformations that do not overlay when the full polypeptide is considered (rmsd = 7.412 Å, Supp. Table S1, Fig. 1C). Yet, the folding of each of the two domains is conserved with the NTD, CTD and linker domains overlaying with rmsd values of 0.536, 0.870 and 4.569 Å, respectively (Supp. Table S1). In chain A, however, the first helix of the CTD (α6) is a long uninterrupted helix ranging from residues 110 to 136, while in chain B this helix is fragmented into two shorter helices (α6_a_ and α6_b_) separated by a 6-residue coil that positions α6_b_ at a 45° angle relative to α6_a_ helix, causing the CTD to adopt a very different orientation relative to the NTD. This disruption of the α6 helix in chain B is essential to accommodate the constraints of the domain-swapping. Moreover, in chain B, the CTD undergoes a further 90° rotation along the longitudinal axis of α6_b_ that positions the helical bundle on the opposite side of the α6 helix compared to chain A (Fig. 1C) and thereby allows chain B to wrap tightly around chain A, making contacts via the NTD, the linker region and the CTD (Fig. 1B). As a result, the two monomers adopt very distinct conformations and this asymmetry creates a marked difference in the two faces of the dimer (Fig. 1B).

A dimer of dimers was also observed by crystallographic symmetry in which two dimers face each other at a 93° angle (Supp. Fig. S2), thereby forming a putative tetramer, with a dimer-dimer interface covering 1044 Å^2^, with 10 H-bonds and 10 salt bridges. We used size-exclusion chromatography coupled to multi-angle laser light scattering (SEC-MALLS) and analytical ultracentrifugation (AUC) to further characterize the quaternary structure of DdrC. Both techniques revealed that a large majority (>90%) of DdrC protein is in the form of dimers with a mass around 45 kDa (Fig. 1D and E). These measurements are in agreement with earlier chemical crosslinking studies that indicated that DdrC could indeed form dimers (29). No tetramers of DdrC were detected by SEC-MALLS and AUC, suggesting that the observed tetramers most likely result from crystal packing. The biological unit thus appears to be the domain-swapped homo-dimer observed in our crystal structure.

All-atom MD simulations of the DdrC homo-dimer confirmed that the dimer was stable throughout the simulation and that the asymmetry of the dimer was also maintained, indicating that this asymmetry observed in our crystal structure does not result from crystal contacts (Supp. Fig. S3). The CTD region, with the exception of the loop linking helices α7 and α8, and the dimer interface of DdrC are particularly stable. In the NTD, the loops and the wHTH motif exhibit some flexibility. Significant movements of the NTD with respect to the CTD were also observed allowing the wHTH of one monomer to come very close and even interact with the C-terminal four-helix bundle of the second monomer (Supp. Fig. S3).

### DdrC-NTD contains a negatively charged wHTH motif

To gain insight into the potential function of DdrC, we performed a search for structural homologues of DdrC using the DALI server. The four-helix bundle in the CTD of DdrC is a very common structural motif found in diverse protein families and is thus not indicative of a particular function. The wHTH motif found in the NTD, on the other hand, has been identified as a DNA-binding motif in several proteins (71–73). The classic wHTH is a positively charged HTH motif followed by a β-hairpin, the “wing”, and preceded by a short β-strand. The conserved motif is usually folded as “β-α-’turn’-α-β-‘wing’-β”. In most wHTH proteins, additional α-helices are packed next to the wHTH motif, usually preceding it in the sequence. In the usual DNA binding mechanism, the HTH part is inserted into the major groove of DNA while the “wing” of the β-hairpin inserts into the minor groove.

The NTD of DdrC exhibits a classic wHTH motif (β3-α1-α2-β4-β5), although the additional α-helix (α3) is located downstream of the wHTH motif in the sequence (Fig. 2A). The wHTH motif of DdrC is also preceded by a hairpin structure composed of β1 and β2. Surprisingly, the electrostatic surface potential of the NTD, calculated with the APBS program (45), indicates that the surface is mainly negatively charged, which would likely prevent DNA binding to this motif (Fig. 2B). A DALI search with the NTD alone confirmed the structural homology of DdrC NTD with other wHTH-containing proteins. The proteins with the highest Z-scores were the human Dachshund protein (PDB code 1L8R, Z score 7.0), the Dachshund-homology domain of human SKI protein (SKI-DHD, PDB code 1SBX, Z score 5.5) and *Bacillus subtilis* RacA (BsRacA, PDB code 5I44, Z score 4.9).

**Figure 2.**
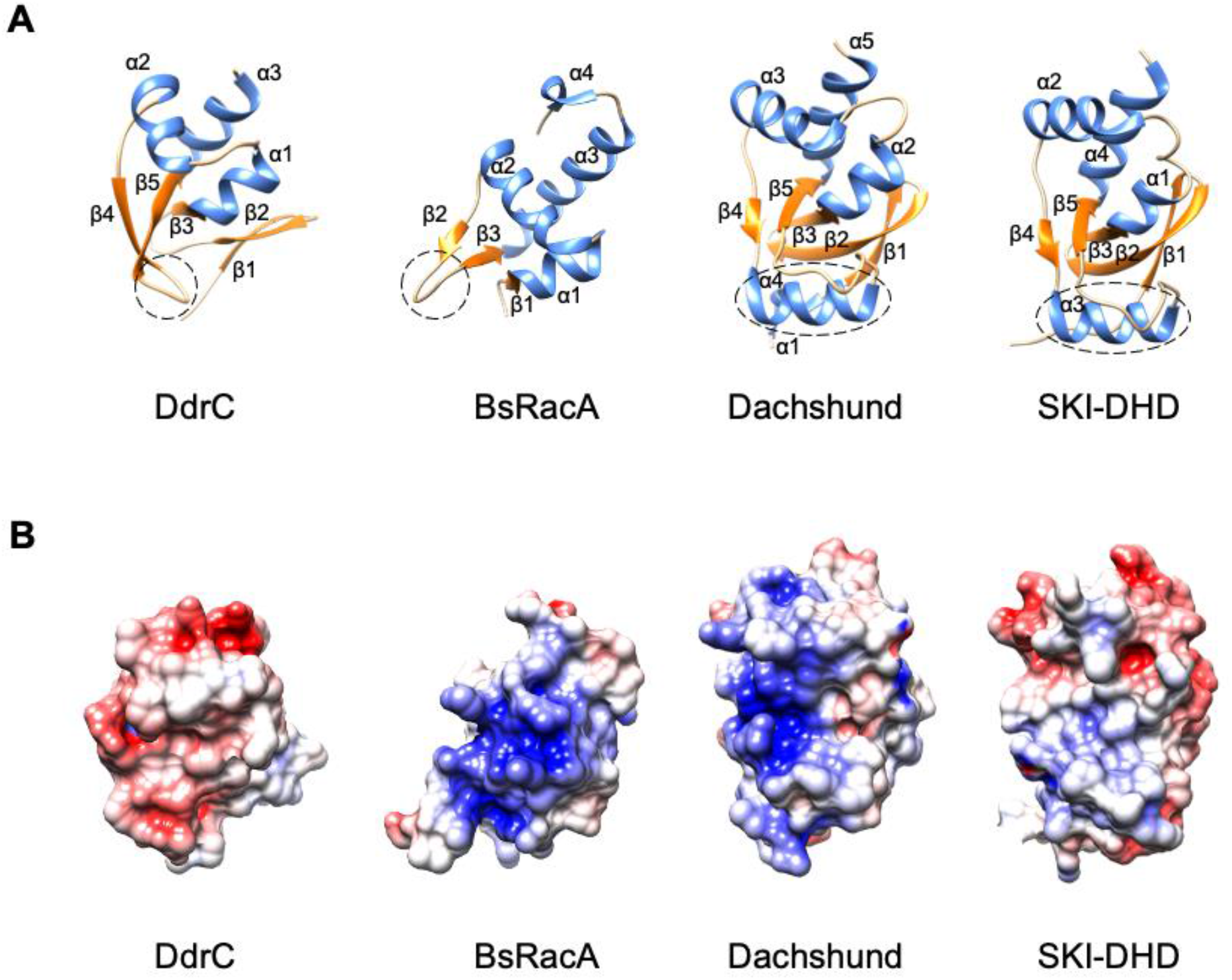
DdrC exhibits a classic yet negatively charged wHTH motif. (A) wHTH motifs of DdrC and structurally similar proteins, BsRacA (PDB code 5I44), Dachshund (PDB code 1L8R) and SKI-DHD (PDB code 1SBX). The proteins are colored based on their secondary structure, with α-helices in blue and β-sheets in orange. (B) DdrC, BsRacA, Dachshund and SKI-DHD are colored by electrostatic surface potential, as calculated by APBS. The color scale is the same for all proteins, ranging from −5 to +5 kT/e, with negative charges in red and positive charges in blue.

The human Dachshund protein (74) and the SKI-DHD domain (75) both of which are involved in transcriptional regulation are very similar to each other and display an unusual wHTH motif, which contains an α-helix inserted in the β-hairpin “wing” between β4 and β5 (β3-α2-α3-β4-**α4**-β5) with the adjacent α-helix located downstream in the sequence (Fig. 2A). As in the case of DdrC, their wHTH motifs are preceded by a β-hairpin structure composed of the two N-terminal β-strands. However, contrary to DdrC, the wHTH motif of the Dachshund protein displays a positively charged surface, which could constitute a DNA-binding interface (74). BsRacA is a kinetochore-like chromosome-anchoring protein that possesses a more classic wHTH motif (β1-α1-α2-β2-β3) and is also positively charged (Fig. 2A and B). The crystal structure of BsRacA in complex with DNA showed that the positively charged wHTH motif is directly involved in DNA binding (76). Unlike Dachshund and BsRacA, the surface of the wHTH motif of SKI-DHD is rather neutral with some electronegative patches (Fig. 2B), and has been proposed to play a role in protein binding rather than in DNA binding (75). These observations suggest that the function of wHTH motifs as DNA-binding sites is more likely associated with their electrostatic surface potential than with their fold. Since the wHTH motif of DdrC exhibits a largely negatively charged surface, it is unlikely to play a direct role in DNA binding.

### DdrC dimer possesses two distinct DNA binding sites

To identify a potential DNA binding site on DdrC, we analyzed the charges displayed at the surface of the DdrC dimer with the APBS program (Fig. 3A). A large positive groove involving mostly arginine residues contributed by the four α-helices of the CTD and α4-α5 of the NTD is present on both sides of the dimer, suggesting that DdrC could possess two distinct DNA binding sites. To test this hypothesis, we performed fluorescence polarization assays with fluorescein-labelled dsDNA oligonucleotides of either 20 (20d5’F) or 50 (50d5’F) base-pairs. DdrC was able to bind efficiently to both DNA substrates (Fig. 3B and C) and in both cases, the best fit was obtained using the two-sites specific binding model (Fig. 3D). For the 20 mer DNA, approximately 50% of the DNA was bound to the high affinity site with a K_D_(Hi) of 60 nM and 50% of the DNA was bound to a second low affinity site with a K_D_(Lo) of 5.42 μM. In contrast, with the longer DNA substrate (50 mer), nearly 75% of the DNA was bound to the high affinity site with a K_D_(Hi) of 130 nM and only a smaller proportion (26.4%) was bound to the lower affinity site with a K_D_(Lo) of 4.97 μM. These data clearly indicate that DdrC dimers possess two distinct DNA binding sites with different affinities for the DNA that can equally accommodate short DNA strands. However, binding of longer DNA fragments to the low affinity site appears to be less favorable.

**Figure 3.**
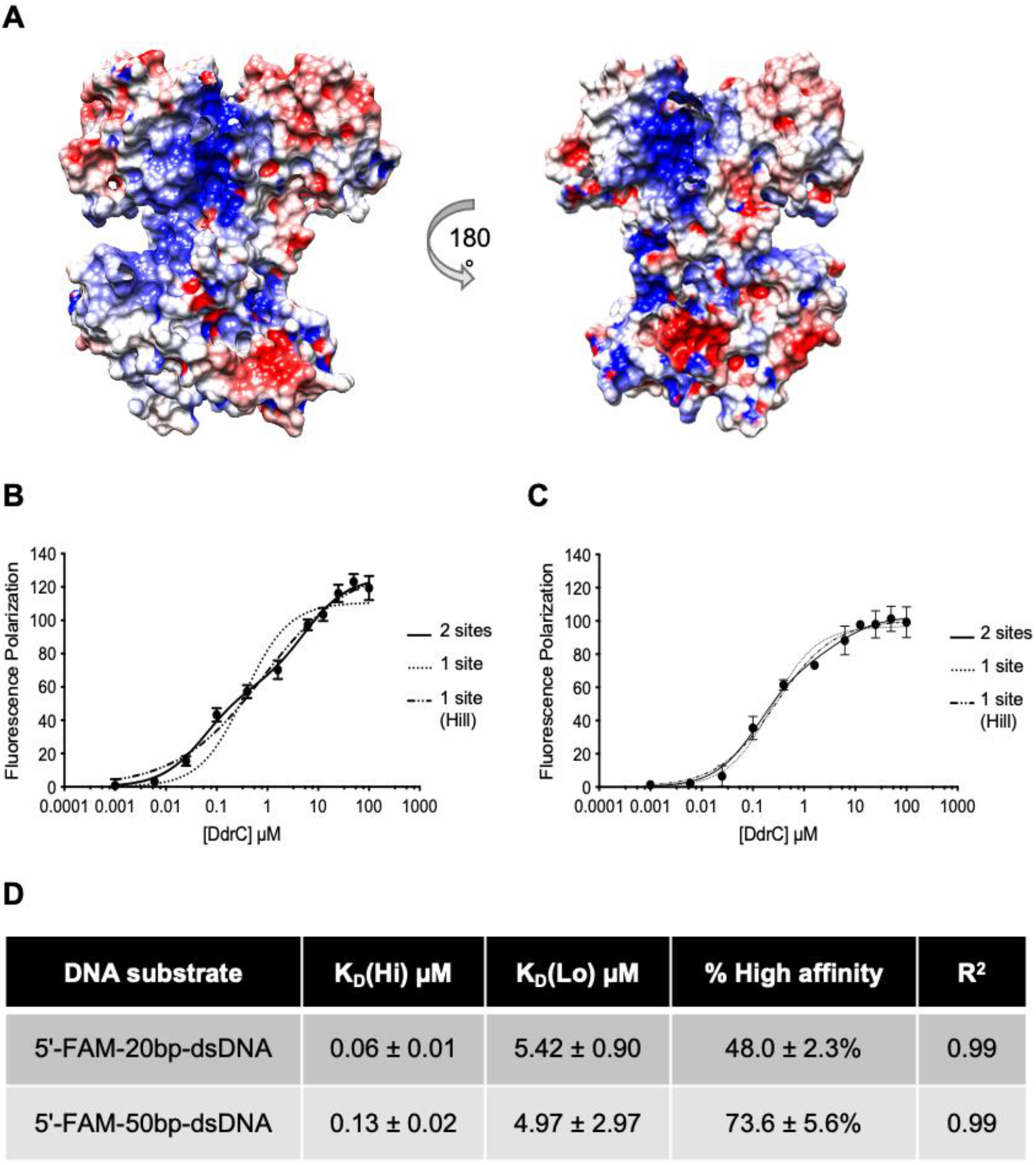
DdrC dimer bears two DNA-binding sites. (A) Depiction of the electrostatic surface potential of the DdrC dimer, as calculated by APBS. Positive and negative charges are colored in blue and red, respectively from −5 to +5 kT/e. (B-C) Fluorescence polarization measurements of 0 to 100 μM DdrC binding to 10 nM 5’-FAM-labelled dsDNA oligonucleotides of 20 bp (B) or 50 bp (C). Mean values with standard deviations of three independent measurements were fitted to one of three models using Prism 8: one-site specific binding (dotted line), one-site specific binding with Hill coefficient (dash-dotted line) and two-sites specific binding (solid line). (D) Binding parameters derived from the fitting of the fluorescence polarization data to a two-sites specific model. K_D_(Hi) and K_D_(Lo) correspond to the binding constants for each of the two sites.

Based on these observations, we built a model in which two 25 bp dsDNA fragments were bound to either face of the DdrC dimer (Fig. 4A). The DNA duplexes were positioned manually along the positively charged grooves of DdrC so as to minimize steric clashes and maintain good geometry. It is interesting to note that the 25 bp dsDNA stretches all the way across these grooves, but adopts a straight conformation on one side and a more bent conformation on the other side of the DdrC dimer where the four-helix bundle of monomer A creates a bulge on the DdrC surface (Fig. 4A). The robustness of this model was then verified by running five independent all-atoms MD simulations over a timescale of 500 ns each (Fig. 4B and Supp. Fig. S4 and S5 and Table S2). As in the case of DdrC alone, only minor changes in the protein conformation were observed during these simulations most of which were restricted to loop regions (Supp. Fig. S4), whereas the two DNA molecules on either side of the DdrC dimer moved substantially to adapt to the protein surface (Supp. Fig. S4 and S5). These movements of the DNA duplex include twisting, bending, sliding along the groove and rotation of the duplex to establish more favorable contacts between the minor grooves of the DNA molecules and the protein.

**Figure 4.**
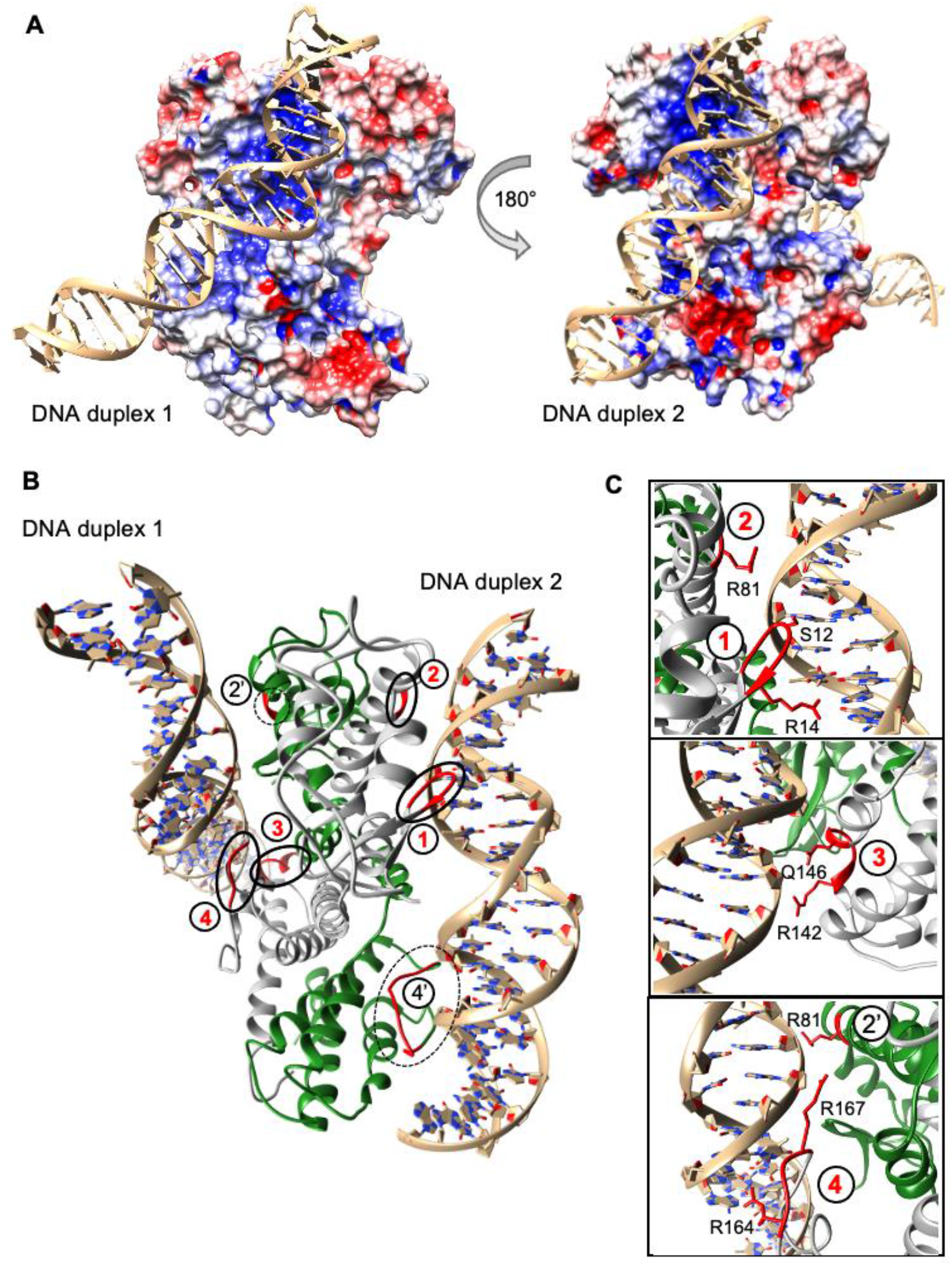
DdrC-dsDNA models derived from MD simulations. (A) Model of DNA-bound DdrC dimer used for MD simulations. Two 25 bp dsDNA fragments were manually positioned along the two positively charged grooves lining each side of the DdrC dimer. (B) Model of DNA-bound DdrC dimer (monomer A in grey and monomer B in green) at the end of MD simulation run3, illustrating the four major contact points (labelled 1-4 in red) and two additional contact points (labelled 2’ and 4’ in black) between the DNA duplexes and the DdrC protein. The regions of DdrC in contact with the DNA are highlighted in red. (C) Close-up views of the major DdrC-DNA contact sites illustrated in (B). The main residues involved in the interactions with the DNA are shown as sticks and are labelled.

Four major contact sites between the DdrC dimer and the DNA duplexes were observed in at least four out of five MD runs (Fig. 4B and 4C, Supp. Table S2 and Fig. S6). Interestingly, the four major contact points are all located in chain A of DdrC that interacts significantly more with the two DNA duplexes than chain B (Supp. Table S2 and Fig. S6). Both the NTD and the CTD domains of DdrC contact the DNA (Fig. 4B and C). The first major contact point involves the N-terminal β-hairpin that precedes the wHTH motif located in the NTD. The second site is located in helix α4 and involves mostly Arg81. The third and fourth contact sites are located in the CTD and involve respectively Arg142 and Gln146 from helix α7 and two positively charged arginine residues (Arg164 and Arg167) situated in the flexible linker between α7 and α8. The second (Arg81) and fourth (Arg164 and Arg167) DNA contact regions were also seen for chain B in at least three out of the five MD runs (Fig. 4B and C; contact points 2’ and 4’). Each face of the DdrC dimer thus contacts a DNA duplex through at least three interaction sites, but as a result of the intrinsic asymmetry of the DdrC dimer, the contact surfaces are quite distinct (Fig. 4B). The interactions between DdrC and DNA are predominantly electrostatic, between the phosphate backbone and positively charged residues, notably arginines, although additional contacts between either protein side chains or the peptide backbone and bases located in the minor groove of the DNA duplexes are also seen (Fig. 4C).

### DdrC alters the topology of plasmid DNA

A previous study based on transmission electron microscopy showed that DdrC was able to condense circular DNA at a high concentration (29). To further investigate the effects of DdrC on plasmid conformation, we incubated supercoiled pUC19 plasmid DNA with increasing concentrations of DdrC and analyzed the resulting DNA-protein complexes by atomic force microscopy (AFM) (Fig. 5 and Supp. Fig. S7). Figure 5 presents representative fields of view obtained at 0, 2, 5, 10 and 20 nM DdrC. To compare the different DNA topologies, we extracted the projected surface areas of individual plasmid-DdrC assemblies (Fig. 5A-E and Supp. Fig. S7) and determined for each field of view the fraction of plasmid molecules that exhibit a condensed conformation (Fig. 5F). At the highest DdrC concentration, almost all the plasmids adopted a highly condensed configuration (93% ± 13%; Fig. 5E and F), which was strikingly different from the 7.5% ± 13% of condensed pUC19sc plasmid molecules in the absence of DdrC (Fig. 5A and F). The fraction of condensed plasmid molecules was clearly seen to increase significantly in a DdrC concentration-dependent manner, suggesting that DdrC may be able to maintain circular plasmid DNA in a condensed conformation.

**Figure 5.**
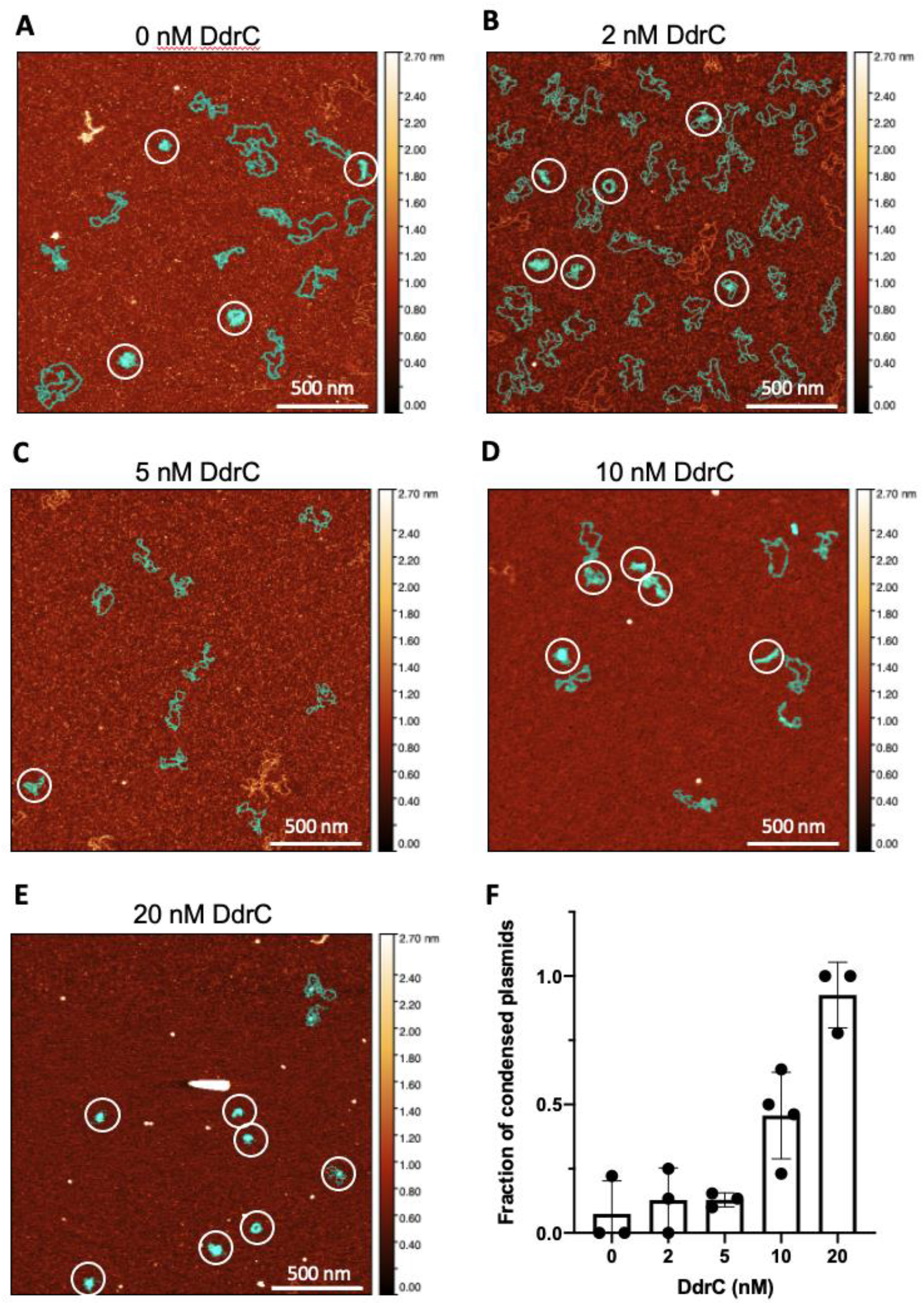
DdrC maintains circular plasmid in a condensed conformation. (A-E) Representative AFM images of 0.5 nM of pUC19sc incubated with 0 (A), 2 (B), 5 (C), 10 (D) and 20 nM (E) DdrC. Additional images are presented in Supp. Fig.S7. All images correspond to 4 μm^2^ areas, in which the assemblies displaying a more condensed conformation are indicated by white circles. The light-blue mask highlights assemblies that have been used in the statistical analysis presented in (F). Assemblies that touch the border of the image or were not clearly identifiable due to unresolved overlapping were excluded from the statistical analysis. The z-scale bar is shown as a color gradient to indicate the distribution of height in the images. Scale bar corresponds to 500 nm. (**F**) Histogram and scatter plot illustrating the mean fraction of condensed pUC19sc-DdrC assemblies as a function of DdrC concentration. The error bars represent the standard deviation of at least three replicates. Individual data points correspond to the fraction of condensed assemblies derived from a single AFM image after estimation of the projected surface area of individual assemblies.

To further explore this property of DdrC, we evaluated whether DdrC could change the topology of circular plasmid DNA by introducing either positive or negative supercoils into relaxed plasmid. For this purpose, we incubated DdrC with a relaxed circular pHOT DNA plasmid, prior to treatment with wheat germ topoisomerase I (TopoI) to relax positive or negative supercoils that might have been introduced by DdrC (Fig. 6). Incubation of TopoI with the relaxed form of the plasmid had no effect on DNA topology in the absence of DdrC (Fig. 6A). In contrast, when the relaxed plasmid was preincubated with DdrC prior to addition of TopoI, several additional topoisomers exhibiting increased supercoiling (faster migration) were observed indicating that DdrC is indeed able to constrain closed circular DNA in a more supercoiled conformation. To distinguish between positive and negative supercoiled topoisomers, plasmid DNA incubated with TopoI alone or with DdrC followed by TopoI were further separated by two-dimensional gel electrophoresis in the presence of chloroquine, a DNA intercalator that unwinds closed circular DNA in the second dimension (Fig. 6B and C). Interestingly, while the starting relaxed pHOT-DNA substrate migrated as slightly positively supercoiled, as expected for relaxed circular plasmid in the presence of chloroquine (77), the incubation of the substrate with increasing concentrations of DdrC generated negatively supercoiled DNA (Fig. 6B and C). DdrC is thus capable of modifying the topology of duplex DNA *in vitro* by generating negative DNA supercoils.

**Figure 6.**
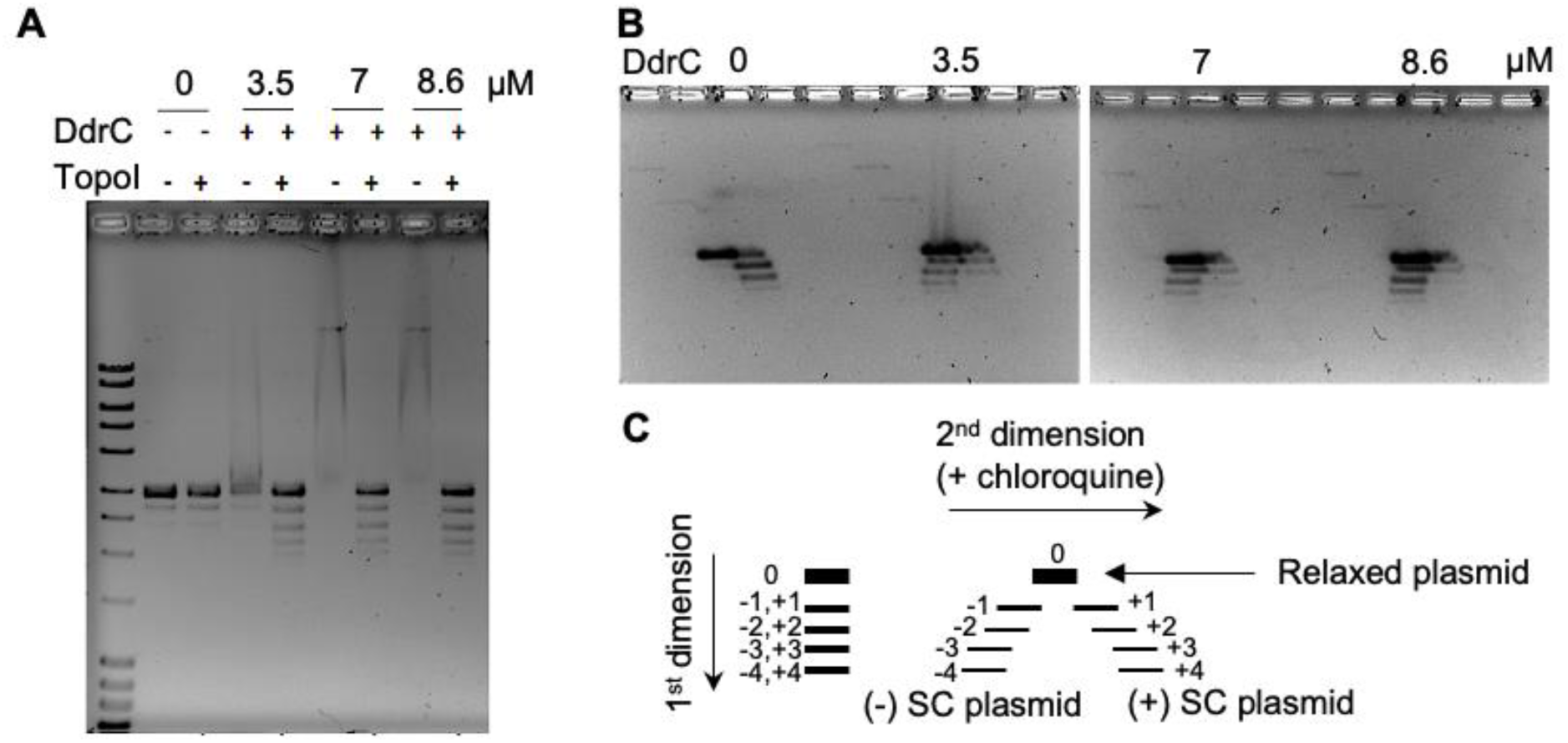
DdrC changes the topology of plasmid DNA by constraining DNA supercoils. (A) Relaxed pHOT plasmid DNA (250 ng) incubated with 0, 3.5, 7 and 8.6 μM DdrC was then treated or not with topoisomerase I (TopoI) from wheat germ. After deproteinization, reaction products were separated by electrophoresis on a 1.2 % agarose gel to resolve topoisomers. Treating relaxed plasmid DNA with TopoI has no effect, whereas treating relaxed plasmid DNA pre-incubated with 3.5-8.6 μM DdrC prior to the TopoI treatment results in a ladder like pattern. (B) The topoisomers resulting from DdrC and TopoI treatment were further separated by bidimensional gel with 3 μg/mL chloroquine included in the gel and buffer in the second dimension. Under these conditions, positively supercoiled ((+)-SC) topoisomers migrated towards the right and negatively supercoiled ((-)-SC) topoisomers towards the left, as illustrated in the schematic diagram shown in (C).

## DISCUSSION

Our crystallographic data reveal that DdrC is composed of two domains, an unusual N-terminal wHTH motif and a more classical four-helix bundle at its C-terminus, which is domain swapped in the DdrC homo-dimer. This domain swapping is facilitated by the rearrangement of a long α-helix, α6 in chain A, into two shorter α-helices, α6_a_ and α6_b_, connected by a 6-residue linker in chain B. This break in the helix creates a highly unusual asymmetric homo-dimer, which was not predicted by current artificial intelligence programs. AlphaFold2 correctly predicted the structure of monomer A, but not of monomer B with the disrupted helix, suggesting that the monomer A conformation is likely more stable. The conformation of monomer B may only be elicited upon protein dimerization. If true, this would mean that interaction of monomer A with monomer B changes the equilibrium conformational energy landscape of monomer B, leading to adoption of a new structure through inducement of a helix break and change in the relative orientation of the two domains — an impressive illustration of structural moonlighting. Alternatively, both structures may exist in solution, even though the conformation of monomer B was not predicted by the machine learning algorithms. Improvements of these algorithms in the future may allow to favor one or the other of the two hypotheses. Regardless, the DdrC dimer structure exemplifies that *de novo* phasing of crystallographic data will in some cases remain the surest pathway towards structure determination.

In DdrC, the domain swapping creates an asymmetric dimer exhibiting two distinct DNA binding surfaces. Our DdrC-DNA models suggest, however, that the conserved wHTH motif that is found in numerous DNA binding proteins is not involved in DNA binding in the case of DdrC (71–73). The electrostatic surface potential of DdrC’s wHTH motif is indeed largely electronegative, precluding a direct involvement in DNA binding. The closest structural homologues of this wHTH motif of DdrC are found in the Dachsund protein (74) and the SKI-DHD domain (75), which also possess unusual wHTH motifs preceded by a N-terminal β-hairpin structure. The Dachshund protein, however, exhibits a positively charged wHTH, while the electrostatic surface potential of the SKI-DHD domain is more similar to that of DdrC and has been shown to be the site of protein-protein interactions (75). The negatively charged wHTH of DdrC may thus also constitute a binding site for a partner protein rather than for DNA.

Taken together, our fluorescence polarization and MD simulations data clearly indicate that DdrC can simultaneously bind two DNA duplexes via its two sides. Six DNA interaction sites were identified on DdrC with elements from both the NTD and the CTD taking part in DNA binding. Interestingly, monomer A contributes significantly more than monomer B to direct contacts with the DNA (four out of the six contact points). The major contact points involve the N-terminal β-hairpin, helices α4 and α7 and the C-terminal flexible linker connecting helices α7 and α8. Although each side of DdrC contacts the DNA in three different regions, the two DNA binding surfaces of DdrC are remarkably different and may well represent the high- and the low-affinity binding sites identified in our fluorescence polarization experiments.

These findings indicate that DdrC may function by bridging DNA duplexes bound on either side of the dimer. This could explain its previously reported DNA circularization and single-strand annealing activities (29), but also its ability to maintain circular plasmid DNA in a condensed conformation as shown in our AFM images. This may be achieved in part at least by neutralizing the negatively charged DNA backbone to allow the close packing of DNA duplexes. DNA compaction by DdrC may also be facilitated by its ability to modify the topology of circular DNA as revealed by our experiments with DdrC coupled to TopoI relaxation activity. DdrC can indeed modify the topology of DNA *in vitro* by constraining DNA supercoils that are subsequently transformed into negative supercoils by TopoI in our assay. This additional function of DdrC may be needed for the reorganization of the nucleoid in response to genotoxic stress.

These features of DdrC are reminiscent of those of NAPs, which play key roles in the organization and tight packaging of genomic DNA in bacterial cells through DNA bending, wrapping and bridging (78–80). The genome of *D. radiodurans* only encodes for a small number of NAPs, with HU and the DNA gyrase complex being the most abundant NAPs associated with *Deinococcus* nucleoids (22, 81, 82). Unlike other bacterial species, the genome of *D. radiodurans* does not encode for a classical DNA bridging NAP such as the nucleoid-structuring protein H-NS. Under normal growth conditions, the HU and DNA gyrase are thus largely responsible for maintaining the high level of compaction of *D. radiodurans* nucleoids, whilst providing sufficient plasticity to allow for the necessary rearrangements associated with cellular activity and cell cycle progression (5). Interestingly, fluorescence microscopy studies have revealed that exposure of *D. radiodurans* to high doses of γ-irradiation induces increased nucleoid compaction (29, 83). Since DdrC is rapidly recruited to the nucleoid following irradiation, we propose that DdrC may function as a DNA damage-induced NAP that contributes to the enhanced level of compaction of the nucleoid after irradiation by bridging DNA duplexes, thereby limiting the dispersion of the fragmented genome immediately after irradiation to facilitate subsequent DNA repair. The DNA gyrase is also over-expressed after irradiation, and may thus also contribute to the increased nucleoid compaction observed following irradiation by modulating the extent of supercoiling of the genomic DNA, a function that may be further enhanced by the binding of DdrC to DNA and its ability to constrain DNA supercoils.

Three hours post-irradiation, once the DNA repair process is almost complete (84), the abundance of DdrC decreases and the cellular distribution of DdrC changes drastically (29). DdrC which was so far evenly distributed throughout the nucleoid relocalizes to foci located near the closing septum between two *D. radiodurans* cells (29). This site corresponds to the location of the *Ter* regions of the chromosomes, where final chromosome segregation occurs, including DNA decatenation of replicated chromosomes (5, 83). At this stage, the nucleoids also progressively recover their original less compacted conformation, perhaps as a result of the changes in the abundance and distribution of DdrC. This intriguing relocalization of DdrC suggests that DdrC may play a second, distinct function at the late stages of the response to DNA damage to ensure that chromosome segregation and cell division do not resume before DNA repair is complete (5, 83). Further studies will be needed to explore the molecular mechanisms underlying this second putative role of DdrC in the response of *D. radiodurans* to severe radiation-induced DNA damage.

## DATA AVAILABILITY

Atomic coordinates and structure factors for the reported crystal structures have been deposited with the Protein Data bank under accession number XXXX.

## SUPPLEMENTARY DATA

Supplementary Data are available at NAR online.

## ACKNOWLEDGEMENT

IBS acknowledges integration into the Interdisciplinary Research Institute of Grenoble (IRIG, CEA). This work used the platforms of the Grenoble Instruct-ERIC center (ISBG; UMS 3518 CNRS-CEA-UGA-EMBL) within the Grenoble Partnership for Structural Biology (PSB), supported by FRISBI (ANR-10-INBS-0005-02) and GRAL, financed within the University Grenoble Alpes graduate school (Ecoles Universitaires de Recherche) CBH-EUR-GS (ANR-17-EURE-0003). We thank Christine Ebel for access to the AUC and SEC-MALLS platforms. This work acknowledges the AFM platform at the IBS. All the calculations were performed on the local LPCT computing cluster.

