## Supplementary Material for "Structural and functional characterization of DdrC, a novel DNA damage-induced nucleoid associated protein involved in DNA compaction"

### Tables S1 and S2
### Supplementary Figures S1-S7

**Table S1:** Comparison of the *de novo* DdrC crystal structure with the top five predicted DdrC models obtained with RosettaFold and AlphaFold2. The table lists the rmsd values for the overlays of either the whole structure or separate domains (NTD, residues 1-97; Linker, residues 98-138; CTD, residues 139-231).

| Alignment to chain A | | | | | | | | | | | |
|---|---|---|---|---|---|---|---|---|---|---|---|
| Rmsd (Å) | X-ray structure | RosettaFold | | | | | AlphaFold2 | | | | |
| | Chain B | Model 1 | Model 2 | Model 3 | Model 4 | Model 5 | Model 1 | Model 2 | Model 3 | Model 4 | Model 5 |
| all atoms | 7.412 | 7.677 | 7.692 | 7.799 | 7.622 | 7.676 | 2.439 | 2.439 | 2.439 | 2.439 | 2.439 |
| residues 1-97 | 0.536 | 2.430 | 2.555 | 2.656 | 2.559 | 2.954 | 0.813 | 0.856 | 0.907 | 0.872 | 0.874 |
| residues 98-138 | 4.569 | 1.858 | 2.101 | 2.348 | 2.093 | 2.414 | 1.084 | 1.129 | 1.096 | 1.092 | 1.229 |
| residues 139-231 | 0.870 | 0.807 | 0.762 | 0.761 | 0.792 | 0.796 | 0.736 | 0.740 | 0.783 | 0.757 | 0.661 |
| **Alignment to chain B** | | | | | | | | | | | |
| Rmsd (Å) | X-ray structure | RosettaFold | | | | | AlphaFold2 | | | | |
| | Chain A | Model 1 | Model 2 | Model 3 | Model 4 | Model 5 | Model 1 | Model 2 | Model 3 | Model 4 | Model 5 |
| all atoms | 7.412 | 11.231 | 11.231 | 11.231 | 11.231 | 11.231 | 6.870 | 6.870 | 6.870 | 6.870 | 6.870 |
| residues 1-97 | 0.536 | 2.467 | 2.682 | 2.730 | 2.488 | 3.439 | 0.881 | 0.936 | 0.998 | 0.984 | 0.973 |
| residues 98-138 | 4.569 | 4.151 | 4.200 | 4.433 | 4.061 | 4.451 | 4.282 | 4.430 | 4.270 | 4.272 | 4.272 |
| residues 139-231 | 0.870 | 1.065 | 1.040 | 1.145 | 0.959 | 1.100 | 0.767 | 0.781 | 0.722 | 0.719 | 0.724 |

**Table S2:** List of residues of DdrC that contact DNA in at least 3 of the five independent runs of MD simulations (500 ns).

| Residue | Chain ID | Min. dist. ± SD (Å) | Mean dist. ± SD (Å) | Run1 | Run2 | Run3 | Run4 | Run5 |
|---|---|---|---|---|---|---|---|---|
| S12 | A | 2.29 ± 0.16 | 4.01 ± 0.56 | X | X | X | X | X |
| R81 | A | 2.37 ± 0.11 | 3.59 ± 0.24 | X | X | X | X | X |
| R167 | A | 2.40 ± 0.12 | 5.24 ± 1.94 | X | X | X | X | X |
| R14 | A | 2.42 ± 0.12 | 5.05 ± 0.73 | X | X | X | | X |
| R81 | B | 2.40 ± 0.21 | 4.48 ± 0.89 | | X | X | X | X |
| Q103 | A | 2.32 ± 0.49 | 4.80 ± 0.85 | X | X | X | | X |
| R142 | A | 2.56 ± 0.05 | 4.28 ± 0.69 | | X | X | X | X |
| Q146 | A | 2.53 ± 0.10 | 3.82 ± 0.38 | | X | X | X | X |
| R164 | A | 2.40 ± 0.09 | 5.56 ± 0.96 | X | X | X | | X |
| R164 | B | 2.56 ± 0.11 | 5.67 ± 1.00 | X | | | X | X |
| R167 | B | 2.43 ± 0.21 | 4.24 ± 0.88 | X | | X | | X |
| R173 | B | 2.48 ± 0.13 | 4.33 ± 0.75 | | | X | X | X |

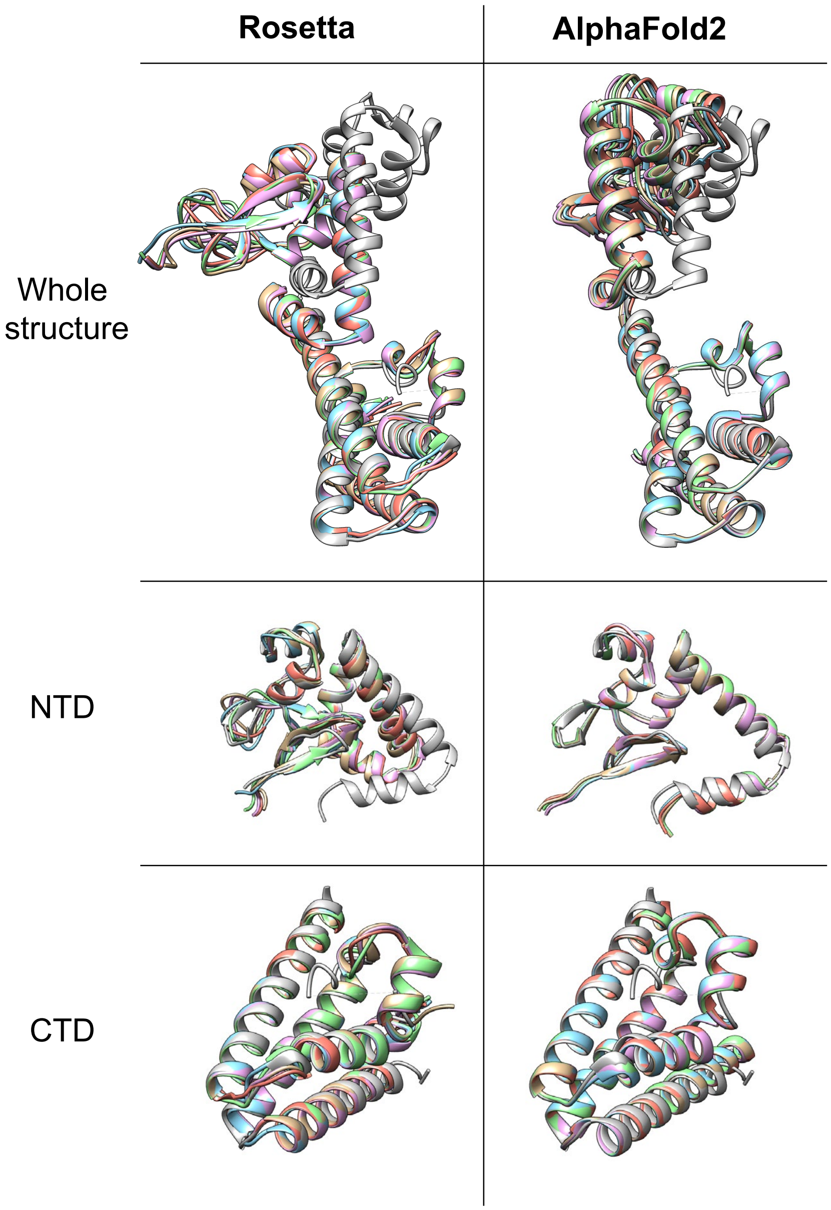


**Figure S1: Overlay of DdrC crystal structure and predicted DdrC models.** Overlay of the NTD (middle) and CTD (bottom) regions of DdrC and overlay of the entire structures (top) after fitting the CTDs. The top five predicted models of DdrC obtained with RosettaFold (left) and with AlphaFold2 (right) are depicted in color (gold, blue, pink, green and red), while the monomer A of the crystal structure of DdrC is colored grey. The CTDs were correctly predicted in both cases, while the NTD was only correctly predicted with AlphaFold2. Nonetheless, the relative orientation of the NTD with respect to the CTD was incorrect in both cases as can be seen in the overlay of the whole structure.

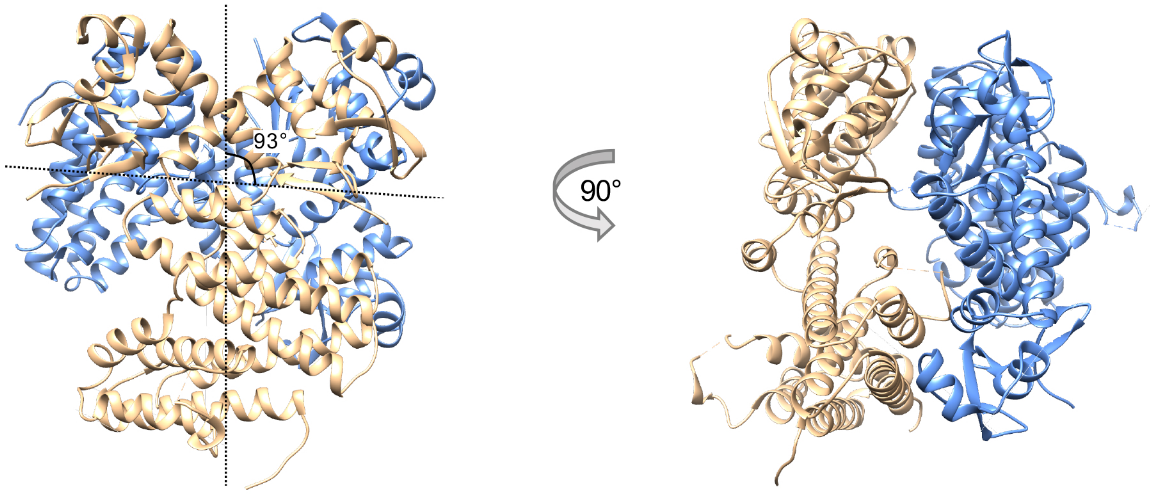


**Figure S2: View of the crystallographic tetramer interface.** Front and side views of a DdrC tetramer, with one dimer colored in beige and the other dimer colored in blue. In the crystal, DdrC forms a tetramer in which one of the dimers is rotated 93° with respect to the other, so that both the chain A and B of each dimer are involved in the interaction.

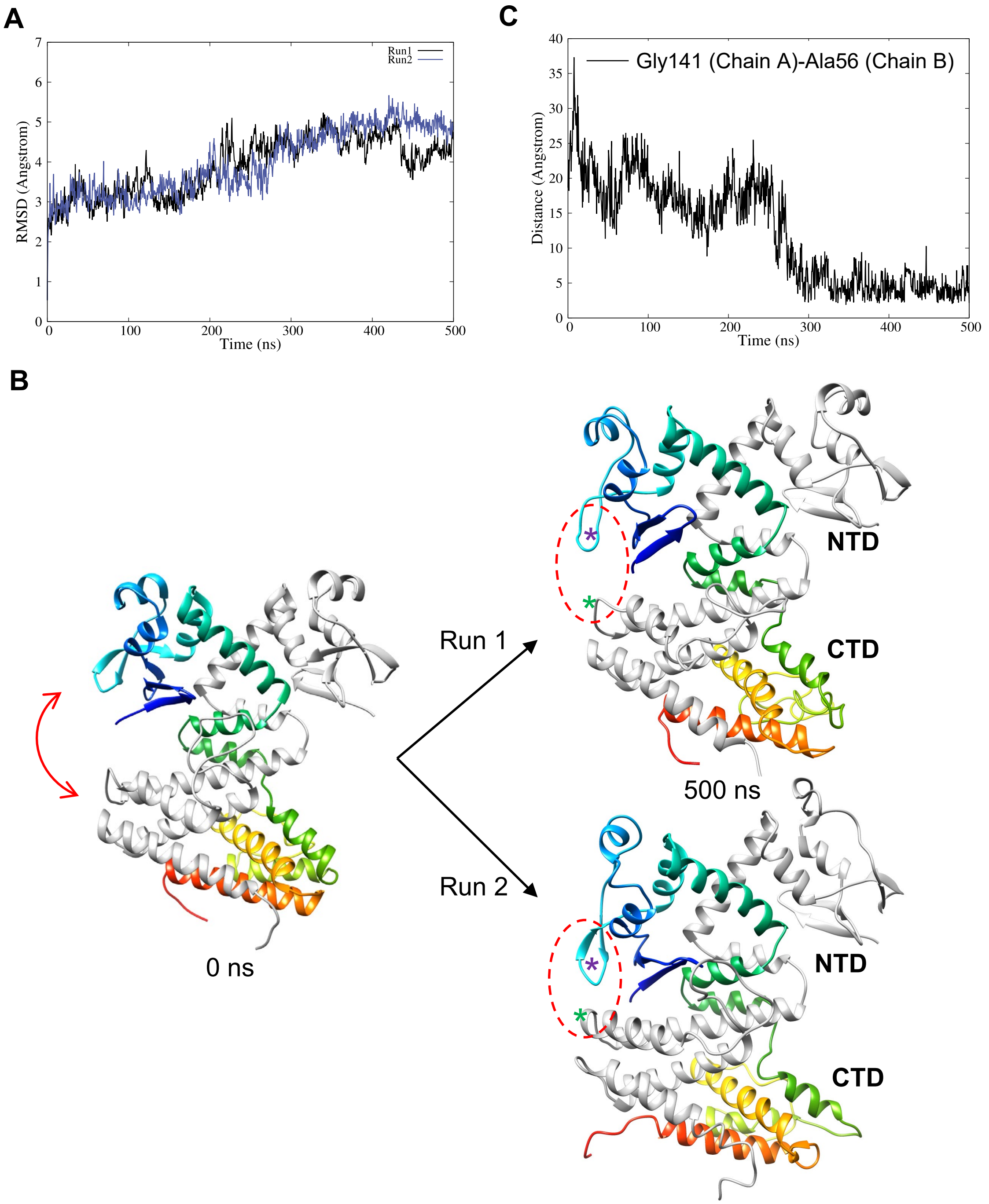


**Figure S3: Molecular dynamics (MD) simulations of DdrC alone (A)** Time series of the rmsd of DdrC extracted from two 500 ns MD simulation runs of DdrC dimer alone. (**B**) Comparison of the conformations of dimeric DdrC at the start of the MD simulations (left) and at the end of the two 500ns runs (right). The dimer is very stable as are all the secondary structure elements, but movements between the NTD and CTD domains (red arrow) are observed, bringing the N-terminal wHTH of monomer B (colored in rainbow colors) in close proximity to the C-terminal four-helix bundle of the monomer A (grey). A red circle with a dashed line indicates this flexible region of DdrC. (C) Time series of the distance between Ala56 located in the β-hairpin structure of the wHTH motif of DdrC chain B (purple star in (B)) and Gly141 of DdrC chain A (green star in (B)) extracted from the 500 ns MD simulation run 2 of DdrC dimer alone.

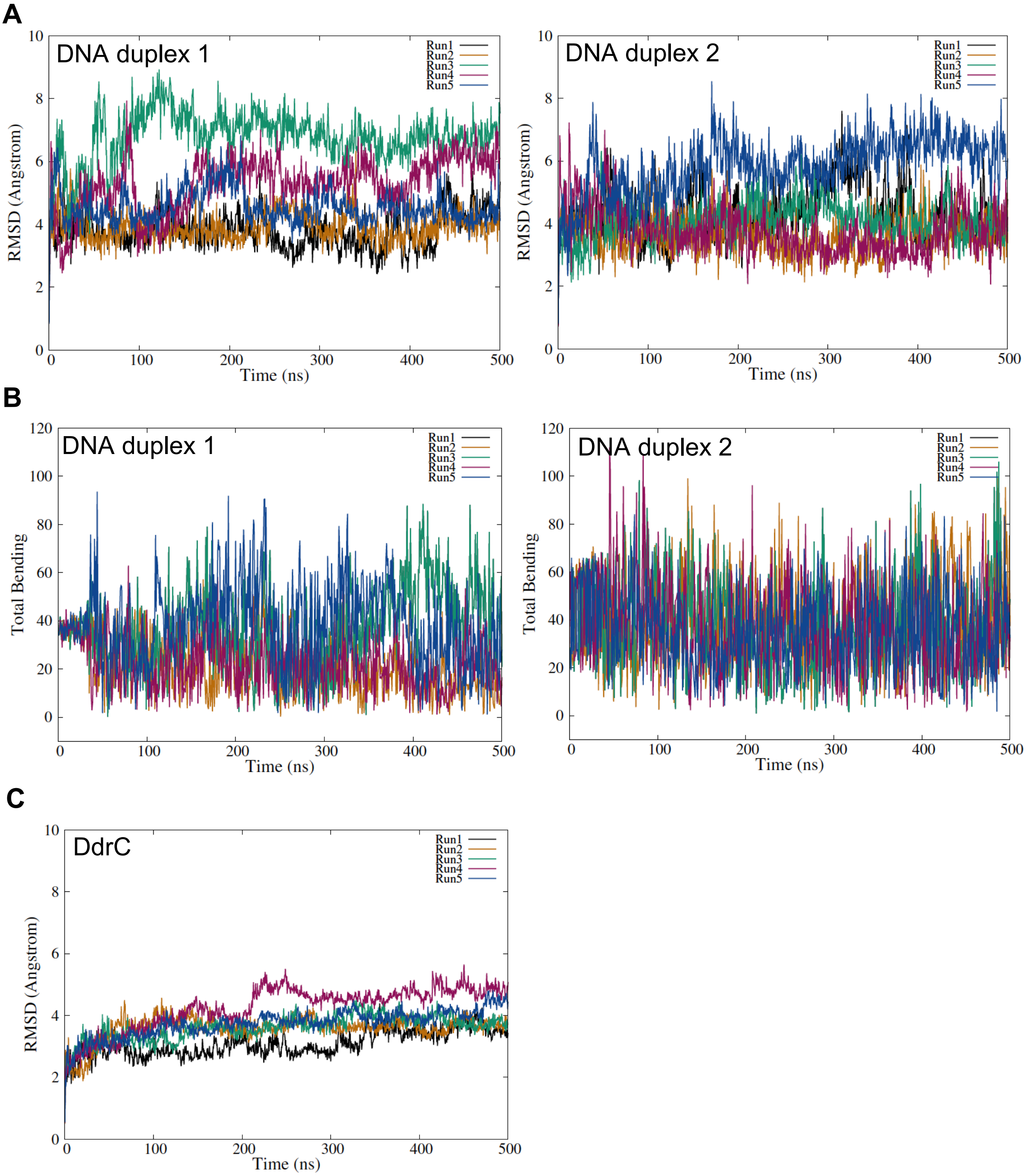


**Figure S4: Molecular dynamics (MD) simulations of DdrC in complex with DNA. (A-B)** Time series of the RMSD (A) and the total bending (B) of the two DNA duplexes (duplex 1, left and duplex 2, right) extracted from the five 500 ns MD simulation runs of DNA-bound DdrC dimer. (**C**) Time series of the RMSD of DdrC extracted from the five 500 ns MD simulation runs of DNA-bound DdrC dimer.

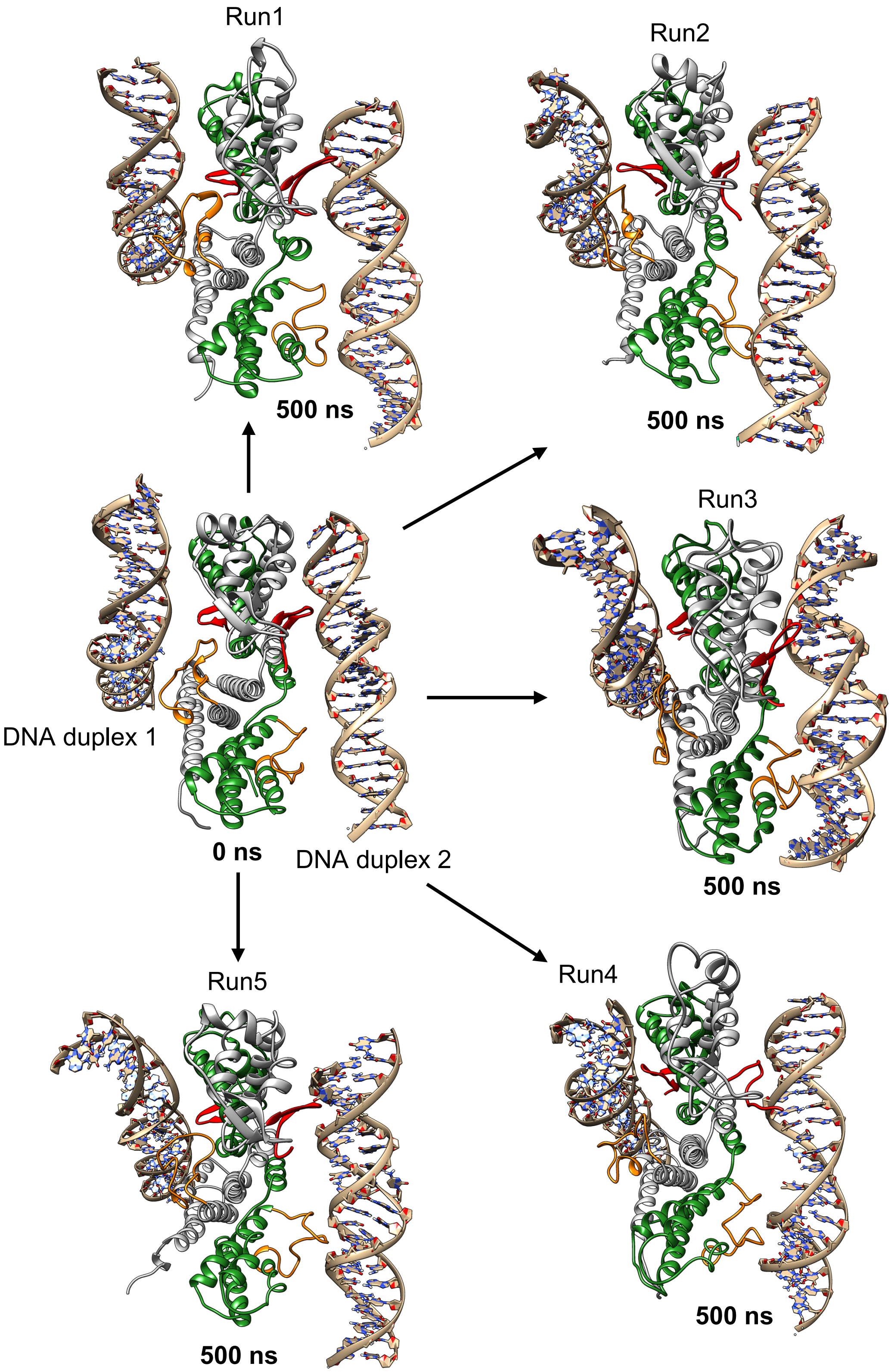


**Figure S5: Molecular dynamics (MD) simulations of DdrC in complex with DNA.** Comparison of the conformations of DNA-bound dimeric DdrC at the start of the MD simulations (left, middle) and at the end of the five independent 500ns runs. The monomers of DdrC are colored in grey (chain A) and green (chain B) and the elements involved in DNA binding are highlighted in red (NTD) and in orange (CTD).

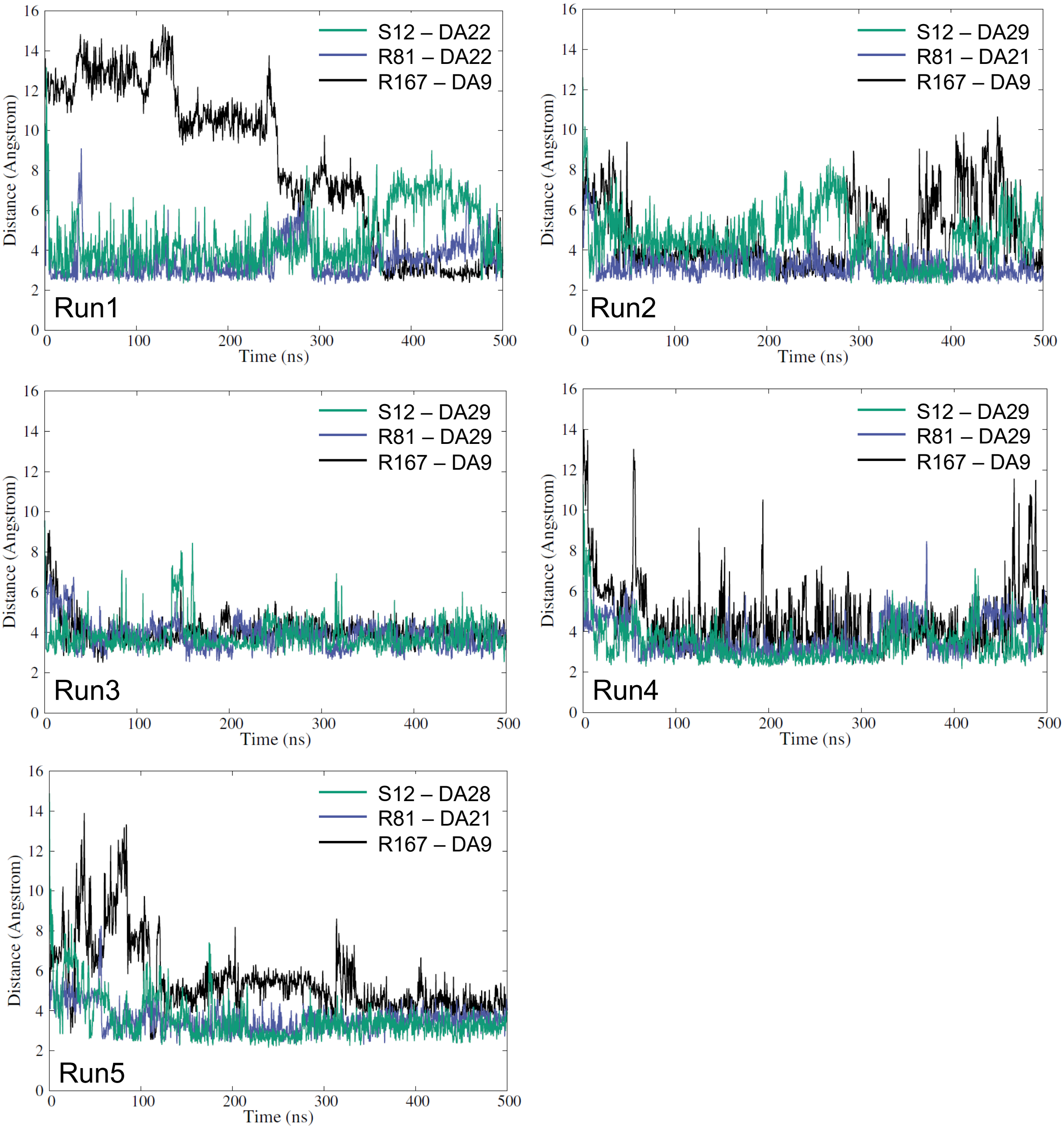


**Figure S6: Evolution of the distances between the three major DNA-interacting residues of DdrC and DNA along the five MD runs.** Time series of the distance between Ser12 (turquoise), Arg81 (blue) and Arg167 (black) of DdrC chain A and the closest DNA atom extracted from the five 500 ns MD simulation runs of DNA-bound DdrC dimer.

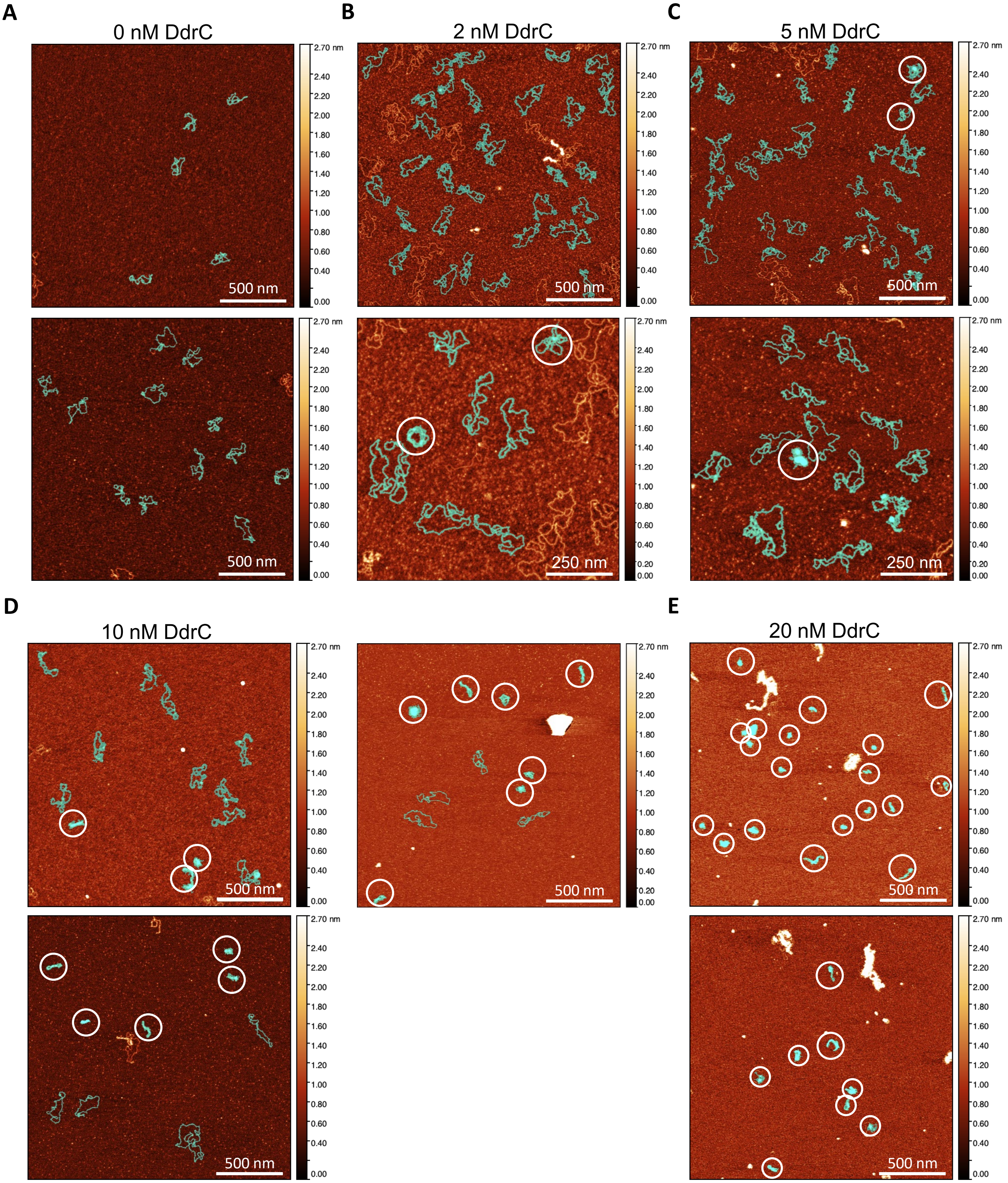


**Figure S7.** DdrC maintains circular plasmid in a condensed conformation. (A-E) AFM images of 0.5 nM of pUC19sc incubated with 0 (A), 2 (B), 5 (C), 10 (D) and 20 nM (E) DdrC. The light-blue mask highlights assemblies that have been used in the statistical analysis presented in Fig. 5F. Assemblies displaying a more condensed conformation are indicated by white circles. The z-scale bar is shown as a color gradient to indicate the distribution of height in the images.
